# A role for heritable transcriptomic variation in maize adaptation to temperate environments

**DOI:** 10.1101/2022.01.28.478212

**Authors:** Guangchao Sun, Huihui Yu, Peng Wang, Martha Lopez Guerrero, Ravi V. Mural, Olivier N. Mizero, Marcin Grzybowski, Baoxing Song, Karin van Dijk, Daniel P. Schachtman, Chi Zhang, James C. Schnable

## Abstract

Transcription bridges genetic information and phenotypes. Here, we evaluated how changes in transcriptional regulation enable maize (*Zea mays*), a crop originally domesticated in the tropics, to adapt to temperate environments. We generated 572 unique RNA-seq datasets from the roots of 340 maize genotypes. Genes involved in core processes such as cell division, chromosome organization and cytoskeleton organization showed lower heritability of gene expression. While genes involved in anti-oxidation activity exhibited higher expression heritability. An expression genome-wide association study (eGWAS) identified 19,602 expression quantitative trait loci (eQTLs) associated with the expression of 11,444 genes. A GWAS for alternative splicing identified 49,897 splicing QTLs (sQTLs) for 7,614 genes. Rare allele burden within genomic intervals with *trans*-eQTLs correlated with extremes of expression in target genes as previously reported for *cis*-eQTLs. Genes harboring both *cis*-eQTLs and *cis*-sQTLs in linkage disequilibrium were disproportionately likely to encode transcription factors or were annotated as responding to one or more stresses. Independent component analysis of gene expression data identified loci regulating co-expression modules involved in phytohormone pathways, cell wall biosynthesis, lipid metabolism and stress response. Several genes involved in cell proliferation, flower development, DNA replication and gene silencing showed lower gene expression variation explained by genetic factors between temperate and tropical maize lines. A GWAS of 27 previously published phenotypes identified several candidate genes overlapping with genomic intervals showing signatures of selection during adaptation to temperate environments. Our results illustrate how maize transcriptional regulatory networks enable changes in transcriptional regulation to adapt to temperate regions.

## Introduction

Whole organism phenotypes are determined by a combination of genetic and environmental factors. Selection scan methods can identify loci with different effects on fitness in natural populations adapted to different environments or different effects on traits deemed desirable by humans in domesticated species^1,2^. However these comparative population genetic approaches typically do not determine the mechanisms by which selected loci alter a given phenotype. Although there are numerous exceptions, genetic variants tend to act on phenotypes by changing the coding sequence and, thus, protein function or by affecting regulatory sequences for transcriptional control, ultimately leading to alterations in protein abundance.

The potential effect size of DNA sequence variants on protein function can be predicted through multiple approaches informed by protein structure, amino acid similarity, and/or evolutionary conservation. Predicting the effect of DNA sequence variants on transcript abundance remains far more challenging, although variant partitioning suggests that 50–55% of variance for various phenotypes is explained by non-coding sequence features^3^. As transcript abundance can be profiled across many individuals, variants associated with variation in the abundance of individual mRNA transcripts can be empirically identified across the genome. Individuals carrying rare alleles in *cis*-regulatory regions for a given gene are disproportionately likely to exhibit gene expression levels in the extreme tails of population expression distribution^2^. Many of the earliest quantitative genetic studies of gene expression regulation were conducted in biparental populations of model organisms, including yeast (*Saccharomyces cerevisiae*)^4^, Arabidopsis (*Arabidopsis thaliana*)^5^, and maize (*Zea mays*)^6^, from which two classes of loci were identified when using the transcript abundance of multiple genes^4–6^: 1) *cis*-acting regulatory variants mapping to the gene in question, playing a role in modulating the expression of a single gene; and 2) *trans*-acting regulatory variation mapping elsewhere in the genome, frequently at “hot-spots”^4–6^. Typically, variants acting in *cis* tend to explain more of the total variance in the expression of their target genes than *trans*-acting regulatory variations^5–7^. However, *trans* acting regulatory variants influencing the expression of multiple genes are frequently identified in single genomic intervals, thus forming “hot-spots” potentially corresponding to variation in a transcription factor or other regulators^4,5,8–10^.

Mapping expression quantitative loci (eQTLs) using recombinant inbred line (RIL) has greater statistical power than biparental populations to identify variants with modest-sized effects mapping to regulatory hotspots associated with variation in the expression of many genes^7,8,11^. Since individual maize transcription factors bind to and regulate the expression of many genes^12^, some or all hotspots may represent functional variants genes encoding transcription factors or other regulatory proteins. Importantly, RIL populations typically do not provide sufficient resolution to resolve mapping to individual candidate genes. By contrast, association mapping utilizing historical recombination events can achieve dramatically higher mapping resolution than biparental populations, particularly in maize where linkage disequilibrium (LD) decays much faster than in many other species^13^, although this comes at a cost of requiring phenotypic data from more individuals^14^. Several eQTL studies using natural diversity panels have offered much higher resolution to map eQTLs to one or several candidate genes or regulatory sequences using transcriptome deep sequencing (RNA-seq). Examples include an analysis of a 224 line maize diversity panel under control and water stressed conditions^15^ and an analysis at two stages of kernel development from 282 diverse maize lines^16,17^. eQTL analyses in natural diversity panels frequently suffer from a lower power to discover the most of small to moderate effect variants segregating in the population, making it challenging to identify “hotspots” affecting the expression of multiple target genes.

Microarray based measurements of gene expression were not specifically designed to distinguish between functionally distinct splice isoforms originating from the same gene. Similarly, 3’ mRNA sequencing provides a scalable and cost-effective mechanism to profile expression from large numbers of individuals but cannot quantify alternative splicing^18^. The expression of different splice isoforms has recently been shown to be regulated both in *cis*- and *trans*^19,20^. Variation in alternative splicing has been reported to be associated with yield^21^, development^22^, stress tolerance^23,24^ and climate adaptation^25^ in plants. As technology for quantifying transcript abundance has advanced, more studies have identified genetically controlled variation in gene expression linked to whole plant phenotypic diversity that contributes to local adaptation, as reviewed by Cubillos et al^26^. For example, seed shattering in field mustard (*Brassica rapa*)^27^ and domesticated rice (*Oryza sativa*)^28^ exhibited diversity due to the same allelic variation in local regulatory elements of *REPLUMLESS* (*RPL*) and *Shattering 1 qSH1*, respectively. Furthermore, apical dominance in maize which played a major role in domestication, is due to an upstream transposon insertion of the *Teosinte branched 1* (*TB1*) gene^29^.

Maize is both a major crop and a model for plant genetics and genomics. After being domesticated 10,000 years ago in what is now south central Mexico^30^, cultivated maize spread into the region that is now the southwestern United States. There, the expansion of maize cultivation range stalled for many years, likely as a result of the poor adaptation of tropical maize to the environments found in more temperate latitudes^31^. Once selection produced maize varieties able to thrive in temperate climates, the crop rapidly spread throughout North America, with maize now being cultivated in a wide range of temperate and tropical regions. Temperate maize differs from tropical maize in a range of phenotypes including flowering time and photoperiod sensitivity^32,33^, as well as tolerance to stresses such as cold, low soil nitrogen, and moisture content^34,35^. By exploiting heterosis, Flint elite temperate maize lines have made for good founders of maize germplasms well adapted to high latitudes^1^. Candidate genes showing significant selective sweep signals in this heterotic pool showed different haplotype diversity between the Flint and Dent groups, with the Flint haplotype promoting early flowering time^1^. This phenotypic divergence is predominantly explained by loci with individual small effects^32^. However a few large-effect loci have also been identified for natural variation in temperate adaptation traits including *Vegetative to generative transition 1 (Vgt1)*, a QTL that produces a 4-5 day change in flowering time and results from polymorphism in a regulatory sequence controlling the transcription of *Related to APETALA 2*.*7 (ZmRap2*.*7)*^36^.

Here we profiled gene expression variation in root system by RNA-seq across a 340-line maize diversity panel comprising a large subset of the Buckler-Goodman maize association panel^37^, augmented with additional diverse maize inbreds. We generated and sequenced two or more biological replicates for most of these genotypes, allowing estimation of broad-sense heritability for each annotated and expressed maize gene. We then mapped both expression QTL (eQTLs) and splicing QTL (sQTLs) using a high density set of 12 million single nucleotide polymorphism (SNP) markers. Consistent with previous expression mapping efforts in association populations, our power to identify *trans*-regulatory hotspots via single gene analysis was limited. However, independent component analysis enabled the identification of latent features associated with the expression of multiple genes, making it possible both to identify genomic intervals controlling these latent transcriptional features, and to assign putative functions to the latent features in processes including cell wall biogenesis, plant-pathogen interactions, fatty acid metabolism, and plant hormone biosynthesis. Using 40 unambiguous tropical genotypes and 52 unambiguous temperate genotypes from the association panel, we identified a set of *cis*-regulatory elements and their targets associated with endosperm color, flowering time and upper kernel shape. Importantly, these variants showed significant differences between tropical maize and temperate maize. We also showed that a set of genes with lower expression heritability in temperate maize are also enriched in functions essential for temperate adaptation such as vegetative to reproductive growth transition, cell cycle regulation and gene silencing.

## Results

### Changes in the heritability of gene expression associated with adaptation and breeding to temperate climates

We generated an average of 18 million sequence reads per sample from 572 RNA samples isolated from maize seedling roots grown under normal conditions (see Methods), consisting of 340 distinct maize genotypes, with 219 genotypes replicated two or more times. We identified 19,565 expressed genes to a level of 1 fragments per kilobase of transcript per million mapped reads (FPKM) or above in at least 458 (80%) samples. Employing the subset of genotypes with two or more replicates, we determined that the distribution of estimated broad sense heritability (H^2^; an estimate of total variance explained by differences between genotypes) is centered around 0.38 and shows an average value of 0.405 (Figure 1a). We observed a modest and negative correlation (Spearman’s ρ = −0.09) between average expression and heritability for expressed genes (Figure 1b). Among 984 non-redundant Gene Ontology (GO) terms assigned to between 25 and 499 expressed genes, genes annotated with six GO terms exhibited significantly higher (>10% and one tail t-test p ≥ 0.05) median heritabilities than the median expression heritability of the overall set of expressed genes, while 19 non-redundant GO terms exhibited significantly lower (<10%) median heritabilities (Figure 1c; see Supplemental document 1). Genes annotated with GO terms linked to metabolic processes, response to oxidative stress anti-oxidation, plant cell wall and photosynthesis were among those whose expression in seedling roots tended to be more heritable across the diversity panel tested (Figure 1c). Genes with lower expression heritability included those annotated as being involved in environmental or pathogen responsive activity as well as histone H3K9 methylation and protein kinases (Figure 1c; see Supplemental document 1).

**Figure 1.**
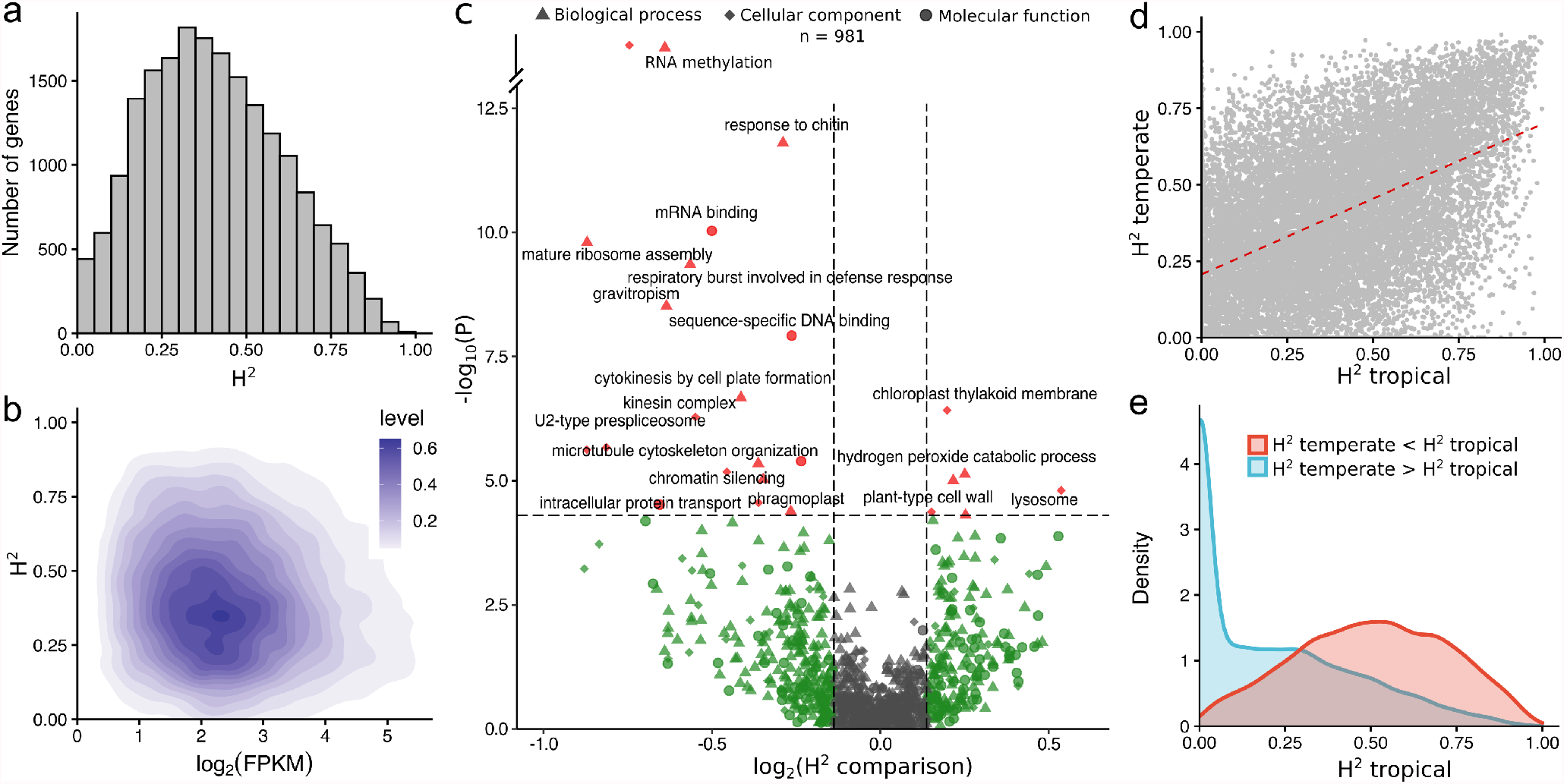
Broad sense heritability calculated from 19565 genes among the 340 maize lines. (**a**) Distribution of broad sense heritability (H^2^); (**b**) Two-dimensional scatter plot between heritability and Log_2_ of mean FPKM values across the population; (**c**) Volcano plot of Gene Ontology (GO) terms that show significant distributional differences across the population. *p*-values are adjusted for a false discovery rate (FDR) ≤ 0.05. (**d**) Distribution of broad sense heritability of genes in the tropical and temperate panels; (**e**) Distribution of tropical expression heritability for genes whose heritability changed in the temperate panel.

The panel profiled here included 40 genotypes of unambiguous tropical origin and 52 unambiguous temperate maize genotypes^38^ (see Supplemental document 2). We noticed a modest but statistically significant (*P* < 1e^−5^; binomial test) skew towards reduced heritability of gene expression among temperate maize lines relative to tropical maize lines (95% confidence interval for linear regression slope coefficient 0.587-0.613) (Figure 1d). In the 52 temperate maize genotypes, the expression heritability of a set of 1,272 genes was less than 20% of heritability in tropical lines. These 1,272 genes exhibited significant enrichment for functional annotations linked to chromosome organization, nitrogen compound metabolism, epigenetic regulation of gene expression, tropism, immune effector process and regulation of flower development (Figure S1; see Supplemental document 3).

### Expression quantitative trait loci reveal the genetic architecture of transcriptional regulation

We mapped expression quantitative trait loci (eQTLs) using MatrixEQTL^39^, implementing an univariate linear model for the expression level of each of the 19,565 expressed maize genes using 12,191,984 SNPs generated from a combination of RNA-seq based SNP calling and data from the maize HapMap3 project (see Methods)^40^. At a threshold *P* < 4.1e-09 (bonferroni correction; 0.05/12,191,984), we identified one or more statistically significant genome-wide association study (GWAS) hits for the expression levels of 11,605 of the 19,565 genes analyzed. We grouped individual significantly associated SNPs into discrete peaks along the maize chromosomes (see Methods) (Figure S2a). We used 11,444 genes with 1 to 10 significant eQTLs for our downstream analyses (these genes are referred to as e-traits below). We excluded 161 genes with more than 10 distinct peaks because they may reflect failure to control for population structure or other confounding factors. We detected single discrete eQTL peak for 7,691 e-traits and for which 7,105 (or 92.4%) co-localized with the position of the annotated gene (a *cis*-eQTL). For the remaining 586 genes (7.6%) the single eQTL peak mapped elsewhere in the maize genome (a *trans*-eQTL) (Figure S2a). Considering e-traits with two or more identified eQTLs (but with only one possible *cis*-eQTL for each e-trait), the overall breakdown was 54.2% of *cis*-eQTLs and 45.8% of *trans*-eQTLs (Figure S2b). In contrast to a previous eQTL mapping study conducted in maize with a bi-parental RIL population^8^, we observed no obvious *trans*-eQTL hotspots (Figure S2c). Overall, of the 11,444 e-traits with at least one significant eQTL peak, 10,618 (92.8%) had a *cis*-eQTL peak (Figure S2d). Consistent with expectations, the cumulative percentage of expression variation explained by all eQTLs identified for a given e-trait was typically lower the estimate of total variance explained from genetic factors (e.g. expression heritability) (Figure 2a, b). *cis*-eQTLs tended to play larger roles in explaining total expression variance for the associated e-trait than *trans*-eQTLs (Figure 2c). *trans*-eQTLs were significantly more likely to represent rare alleles in this maize population (defined here as alleles with a minor allele frequency of <0.2) than *cis*-eQTLs (Figure 2d). Rare alleles at *cis*-eQTLs were shown previously to be associated with extremes in expression^2^. We thus evaluated whether similar associations held true for *trans*-eQTLs with smaller effect size than *cis*-eQTLs. For genes with *trans*-eQTLs, 5 kb regions flanking *trans*-eQTLs showed a greater frequency of rare alleles (allele frequency of <0.2) among genotypes at either extreme of the distribution of expression level for the associated e-trait based on the expression rankings of the 340 lines (Figure 2e, *blue*). We observed no such trend when eQTLs were randomly assigned to e-traits (Figure 2e, *red*). *trans*-eQTLs identified in a population composed of 200 randomly selected genotypes predicted the same association of rare alleles near *trans*-eQTLs in the remaining 140 genotypes (Figure 2f, *blue*). These results are consistent with disregulation of expression induced by *trans* sites imposing similar fitness burdens as those previously observed at *cis*-regulatory sites.

**Figure 2.**
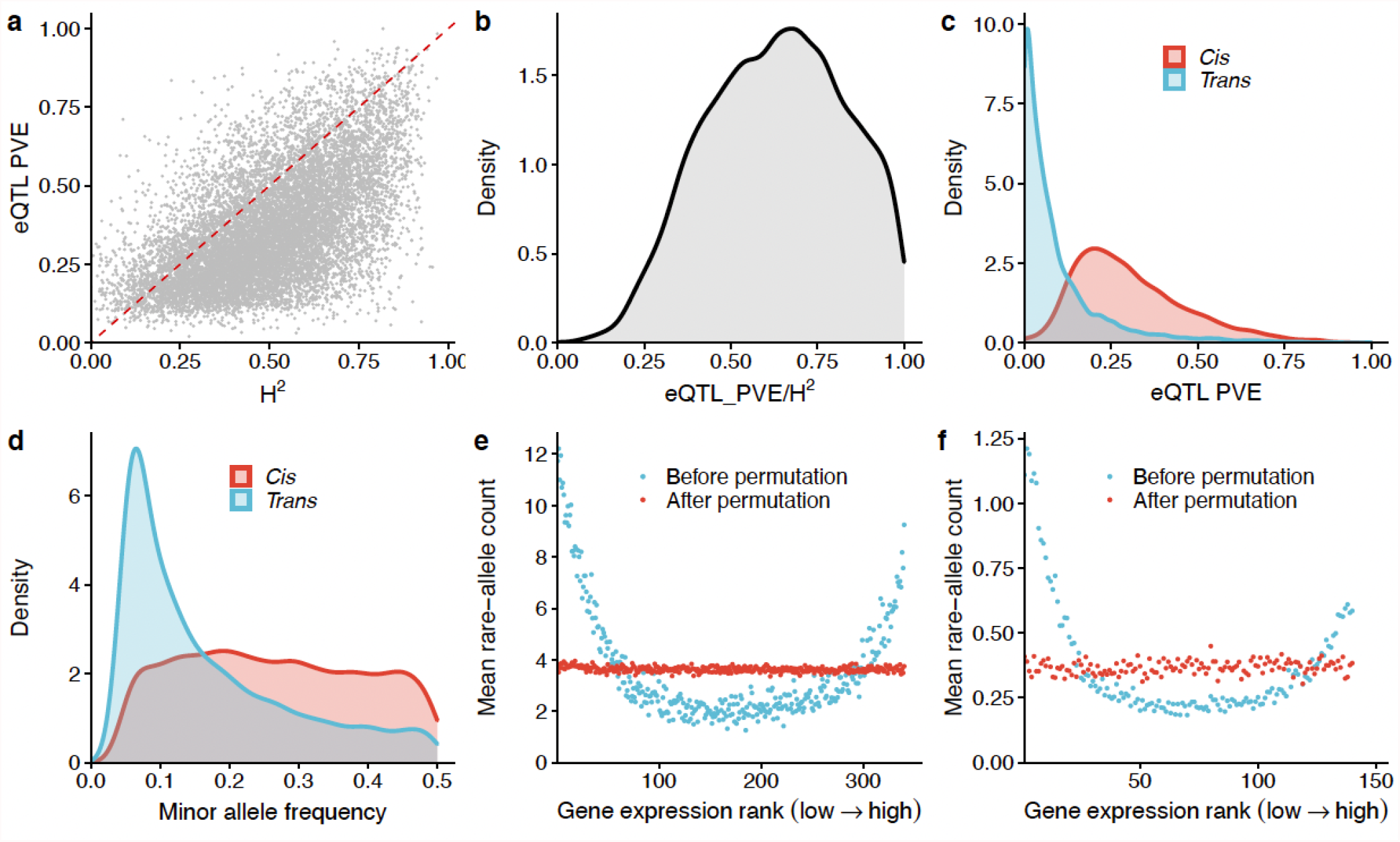
eQTLs and their effects on gene expression. (**a)** Relationship between expression heritability and cumulative percentage of expression variation explained (PVE) for all eQTLs identified for a given gene. (**b**) Expression variation explained by two types of eQTLs. (**c**) Proportion of expression heritability (H^2^) explained by *cis-*and *trans-*eQTLs. (**d**) *trans*-eQTLs are enriched in rare alleles compared to *cis*-eQTLs. (**e**) Rare *trans*-eQTLs are associated with extremes in gene expression, as identified in the entire population. (**f**) *trans*-eQTLs identified in a random subset of 200 genotypes predict the same pattern in the remaining 140 genotypes as observed in (e).

### Alternative splicing QTLs are co-opted with eQTLs for transcriptional regulation

Alternative splicing is a complex regulatory process involved in co-transcriptional and post-transcriptional regulatory mechanisms^42–44^. Variational transcription can be reflected by mRNA levels (whole gene expression) controlled by eQTLs and by the ratio of transcript isoforms mediated by splicing-QTLs (sQTLs). The RNAseq strategy used in this study captured sequence data from all portions of transcripts rather than solely at the their 3’ regions, allowing the quantification of splicing variation across samples and genotypes. For each putative intron, we calculated the percent-spliced-in (PSI) value, defined as the count of exon-exon junction reads normalized by the average per base read coverage of all reads in the intron (see Methods). Based on the reads mapping to intronic regions retrieved from the transcripts assembled by StringTie (v2.1)^45^, we identified 88,888 variable splicing events supported by at least five junction reads in at least 5% of all RNAseq samples with standard errors of PSI≥ 0.01 (see Methods). For each splicing trait (s-trait), we employed the PSI values for splicing-QTL (sQTL) analysis, following the same methodology employed for eQTLs. After excluding s-traits associated with more than 10 distinct sQTL peaks and those associated with annotated genes below the expression threshold, we retained 49,897 sQTLs associated with 16,437 straits in 7,614 annotated genes for downstream analysis (corresponding to two s-traits per gene on average). We detected more *trans*-sQTLs than *cis*-sQTLs (Figure S3b) while *cis*-sQTLs tended to have larger effects and explained a larger proportion of splicing variation than *trans*-sQTLs (Figure S3c), in agreement with a previous report employing gene expression data from developing maize kernels^20^. In contrast, to the pattern observed for *cis*-eQTLs which we frequently observed at the annotated transcription start sites (TSSs) and transcription end sites (TESs) of their associated genes (Figure 3a), *cis*-sQTLs were highly enriched in the gene body (Figure 3a), particularly at the splicing sites of introns ((Figure 3b). We resolved the distance between *cis*-sQTLs and their targets within 1 kb, suggesting that the rapid decay of LD in this maize population enables high resolution mapping of sQTL locations (Figure 2h).

**Figure 3.**
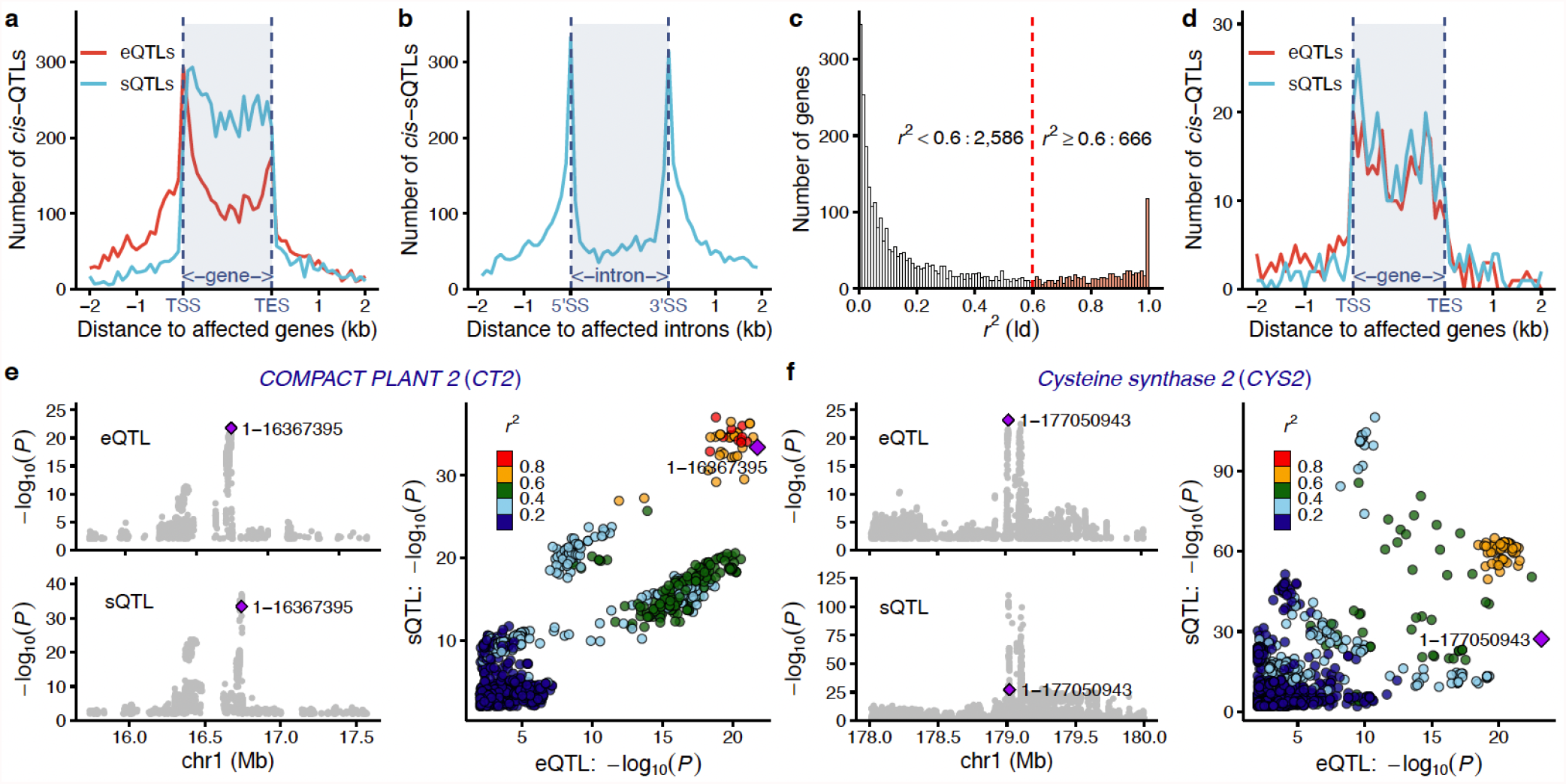
Genes with co-localized *cis*-eQTL and *cis*-sQTLs. (**a**) *cis*-expression QTLs (*cis*-eQTLs) are enriched at transcription start and stop sites, while *cis*-splicing QTLs (*cis*-sQTLs) tend to reside in gene bodies. (**b**) Rapid linkage disequilibrium (LD) decay in the maize genome allows a highly resolved location for *cis*-sQTLs relative to their targets. (**c**) LD (r^2^) distribution of the *cis*-eQTLs and *cis*-sQTLs identified for the same genes. (**d**) Co-localization of highly linked (r^2^ >0.6) *cis*-eQTLs and *cis*-sQTLs relative to their target genes. (**e**) *COMPACT PLANT2* (*CT2*) is co-regulated by *cis* regulatory elements affecting both gene expression levels (eQTL) and splicing variation (sQTL, modified from the output of the locus comparer^41^). (**f**) *Cysteine synthase 2* (*CYS2*) is associated with both a *cis*-eQTL and a *cis*-sQTL, but the two regulatory elements function independently (modified from the output of the locuscomparer^41^).

In 3,252 cases, we determined that an individual maize gene is associated with both a *cis*-eQTL and at least one *cis*-sQTL (Figure 3c). However, in many cases the peak SNPs of the *cis*-eQTL and the *cis*-sQTL were not in LD with each other. To reveal the potential colocalization between *cis*-eQTLs and *cis*-sQTLs, we calculated the R-squared values (*r*^*2*^) for LD between the peak SNP for the *cis*-eQTL and *cis*-sQTL for the same gene, resulting in the identification of 666 genes with colocalizing *cis*-eQTL and *cis*-sQTL peaks (*r*^*2*^ ≥0.6) (Figure 3c). The density of *cis*-eQTLs and *cis*-sQTLs was similar for these 666 genes in gene bodies (Figure 3d). Among different types of splicing variations, 358 out of these 666 genes exhibited variation in intron retention rates, and their expression levels are highly correlated with the splicing level (PSI) of the introns (with a false discovery rate [FDR] <0.05). However, the direction of correlation was not consistent. Indeed, 226 out of these 358 cases exhibited a positive correlation with their expression levels, while the remaining 132 cases showed a negative correlation (chi-square test *p*-value = 6.76e _-7_). This result was consistent with a previous report of the “intron-mediated enhancement” phenomenon whereby the number of introns in a gene tends to be positively associated with expression levels^46^. Mutations in *COMPACT PLANT2* (*CT2*; *Zm00001d027886*) are associated with shorter plant height and wider meristems, leaves, and ears; *CT2* encodes a G*α* subunit of a heterotrimeric GTP binding protein^47^. In the population analyzed here, both overall *CT2* expression and alternative splicing of the first intron (within the 5’ untranslated region) were controlled by one linked *cis*-eQTL and one *cis*-sQTL (*r*^*2*^ = 0.86) (Figure 3e). *Cysteine synthase 2* (*Cys2*; *Zm00001d031136*) provided an example of comparatively unlinked expression and splicing variation targeting the same gene. *Cys2* expression level was associated with a *cis*-eQTL located in the *Cys2* promoter region, while a *cis*-sQTL within the gene body modulated the retention level of the first intron in the 5’ UTR. Despite being located only 7.4 kb apart, the peak SNPs for these *cis*-eQTL and *cis*-sQTL in *Cys2* were only modestly linked with each other (*r*^*2*^ = 0.33), suggesting that these two SNP markers may define separate functional variants controlling expression levels and transcript splicing (Figure 3f).

### Independent component analysis revealed latent *trans* regulation hubs for co-expression modules

eQTL analysis in natural diversity populations frequently lacks the power to discover most of the small to moderate effect variants making it challenging to identify regulatory hotspots. Using the method proposed by Rotival et al^48^, we employed independent component analysis of the expression matrix (572 RNAseq datasets x 19,565 expressed gene models) to generate a signature matrix of latent features reflected in the expression patterns of multiple genes. A set of 164 independent components explained approximately 80% of the total variance of the expression matrix (Figure S4). After filtering independent components based on module distribution kurtosis^49^, we retained 43 independent components associated with the expression of 24 to 743 genes for downstream analysis. Heritability of these 43 independent components varied from 0 to 0.97, as determined by the expression matrix (Figure 4a). Independent components associated with many genes tended to exhibit lower estimated broad sense heritabilities (Figure 4a). Plant reactome^50,51^ enrichment analysis indicated that independent component 149 (IC149) explains variation in the expression of genes associated with cellulose, xylan and tricin biosynthesis; IC32 was associated with jasmonic acid signaling, glycolipid desaturation, salicylic acid signaling, choline biosynthesis III, and phaseic acid biosynthesis, while IC96 was associated with root hair development, jasmonic acid signaling, gibberellic acid signaling, and ethylene biosynthesis from methionine (Figure 4b); Kyoto Encyclopedia of Genes and Genomes (KEGG) enrichment analysis indicated that IC32 explains variation in the expression of genes involved in plant-pathogen interactions and mitogen-activated protein (MAP) kinase signaling pathways; IC46 was associated with fatty acid metabolism; and IC96 was involved in plant-pathogen interactions and plant hormone signal transduction (Figure 4c).

**Figure 4.**
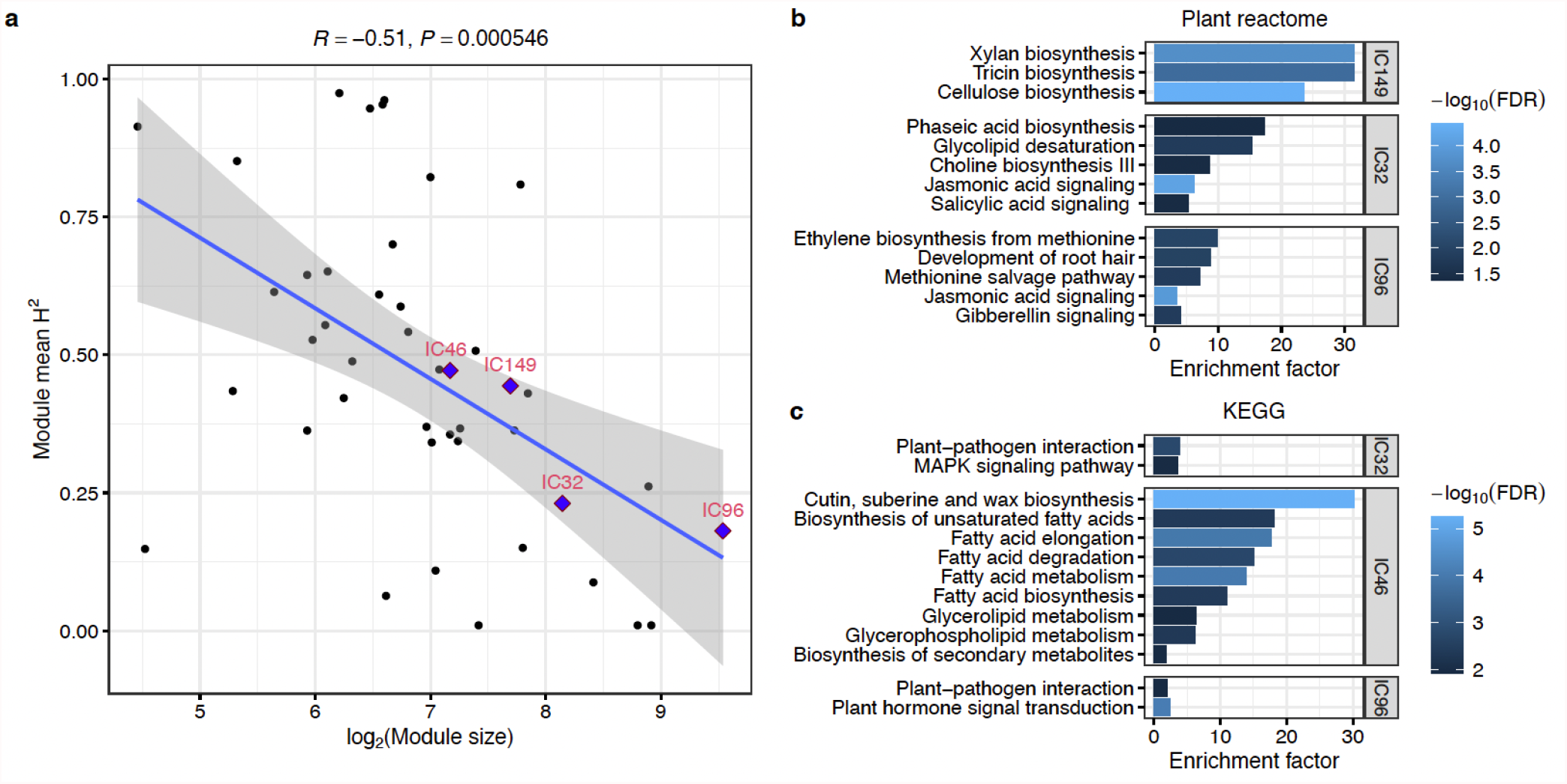
Independent component analysis identified co-expression modules related to important growth and stress response related pathways. (a)The heritability of independent components is negatively correlated with the number of their associated co-expression gene modules. (**b**) Plant reactomes enriched in the co-expression gene modules associated with independent component 149 (IC149), IC32 and IC96. (**c**) Kyoto Encyclopedia of Genes and Genomes (KEGG) term enrichment among the co-expression gene modules associated with IC32, IC46 and IC96. (**b**-**c**) Enrichment factor indicates the fold-change of the percentage of the number of genes associated with the plant reaction in the specific gene modules over that in the background gene set. *p*-values after FDR correction at 0.05 are color-coded.

We conducted a GWAS for each of the independent components, followed by grouping of individually significant trait associated SNPs into distinct peaks as described above. A peak represented by the lead SNP located at 42.3 Mb on chromosome 3 (3:42,334,086) was associated with variation in the expression of 26 out of 207 genes in expression module 149 (IC149) with an enrichment *p*-value of 4.9e^−6^ and was linked to Zm00001d040415 encoding *MIZU-KUSSEI 1* (*MIZ1*) involved in lateral root development by maintaining auxin levels^52–54^, which is in agreement with cellulose and xylan biosynthesis reactions being enriched in the plant reactome for IC149 (Figure 4b). Similary, a peak represented by the lead SNP located at 79.1 Mb on chromosome 4 (4:79,115,907) was associated with variation in the expression of 58 out of 283 genes in expression module IC32 with an enrichment *p*-value of 3.04e^−5^. The SNPs in this peak were strongly linked to a gene encoding trehalose-6-phosphate synthase 9 (ZmTRPS9) whose exprssion and activity is altered in maize *tb1* mutant^55^. Moreover, TRP9 was also involved in disease responses in tomato (*Solanum lycopersicum*) and southern rust^56,57^ in agreement with IC32 being functionally enriched for phytohormone signaling and plant pathogen interactions (Figure 4b,c). A peak represented by the lead SNP located at 281 Mb on chromosome 1 (1:280,872,052) was associated with variation in the expression of 45 out of 743 genes expression module IC96 with an enrichment *p*-value of 1.33e^−5^. The SNPs within this peak are strongly linked to a gene encoding an Acyl-CoA N-acyltransferase with RING/FYVE/PHD-type zinc finger protein (Zm00001d034004) associated with plant height variations in response to drought in maize^58^. While another peak represented by the lead SNP located at 163.2 Mb on chromosome 3 (3:163,218,947) associated with variation in the expression of 23 out of 743 genes, with an enrichment *p*-value of 6.65e^−10^ and is linked with Zm00001d042362, a homolog of Arabidopsis *EXCESS MICROSPOROCYTES1* (EMS1), which encodes a putative leucine-rich repeat receptor protein kinase that controls somatic and reproductive cell fates^59^ (Table S2).

### Genomic mechanisms underlying temperate adaptation

To identify genomic regions exhibiting significant selective sweep signals for adaptation to temperate environment, we performed cross-population composite likelihood ratio (XP-CLR) and fix index (Fst) scan in 100-kb windows with a 10-kb sliding window (see Methods). We obtained 2,503 annotated maize genes located within 100-kb genomic intervals above the 90^th^ percentile for both observed haplotype frequency difference (XP-CLR) and fix index (Fst) in a comparison of 40 unambiguous tropical and 52 temperate maize genotypes included in this study (Figure 6a & b). The highest XP-CLR value identified in this scan was for a genomic interval encompassing a maize homolog of *Arabidopsis PREFOLDIN SUBUNIT-2* (*PFD2*, Zm00001d007490). This gene exhibited a slight change in expression heritability between tropical and temperate populations and a modest change in allele frequency of a peak SNP associated with the *cis*-eQTL detected for this gene (Figure 6 c). The highest Fst value identified was for a genomic interval comprising a maize homolog of Arabidopsis *VERNALIZATION-INSENSITIVE-LIKE PROTEIN −1* (*VIL1*, Zm00001d041715). This gene showed a large decrease in expression heritability between tropical and temperate populations, as well as a large shift in allele frequency for the *cis*-sQTL associated with splicing (Figure 6d). Of the 1,058 potential target genes of selective sweeps between tropical and temperate maize germplasm that are associated with *cis*-eQTLs or *cis*-sQTLs in maize roots 171 showed a greater than 20% reduction in estimated heritability of gene expression between tropical and temperate lines including *Yellow 1* (*Y1*, Zm00001d036345)^60^, a target of artificial selection in temperate maize lines; *VIL1*^61^, the basic helix-loop-helix (bHLH) transcription factor gene *ZmbHLH125* (Zm00001d045212)^62^; as well as additional genes potentially involved in temperate adaptation based on either functional characterization or functional prediction (see Supplemental document 5).

**Figure 5.**
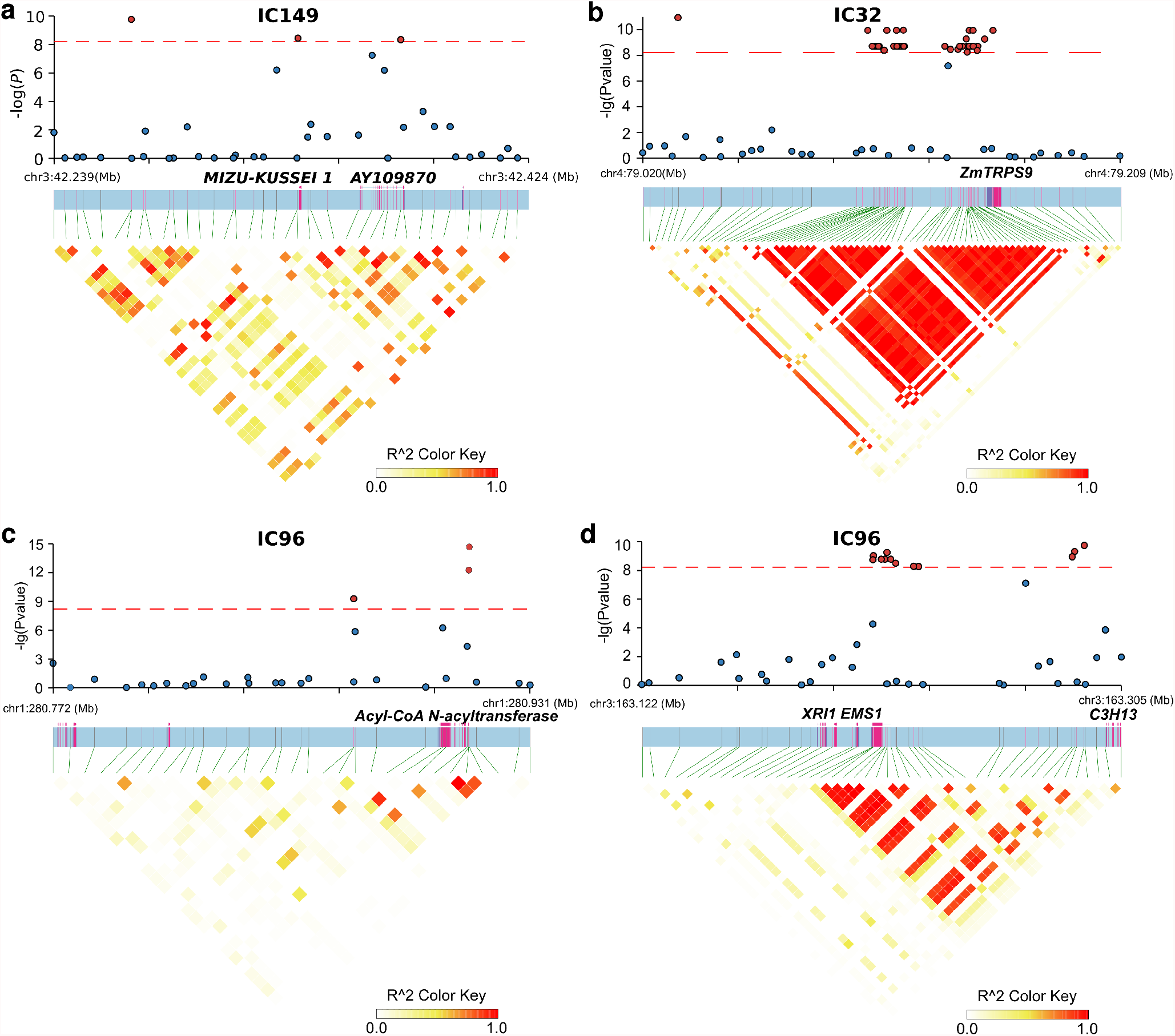
SNP-module association identified candidate global trans-regulators. (a) The significant SNPs associated with IC149 is linked to gene *MIZU-KUSSEI 1*. (**b**) Plant reactomes enriched in the co-expression gene modules associated with IC149, IC32 and IC96 respectively. (**c**) KEGG term enrichment among the co-expression gene modules associated with IC32, IC46 and IC96. (**b**-**c**) Enrichment factor indicates the fold-change of the percentage of the number of genes associated with the plant reaction in the specific gene modules over that in the background gene set. *p*-values after FDR correction at 0.05 are color-coded.

**Figure 6.**
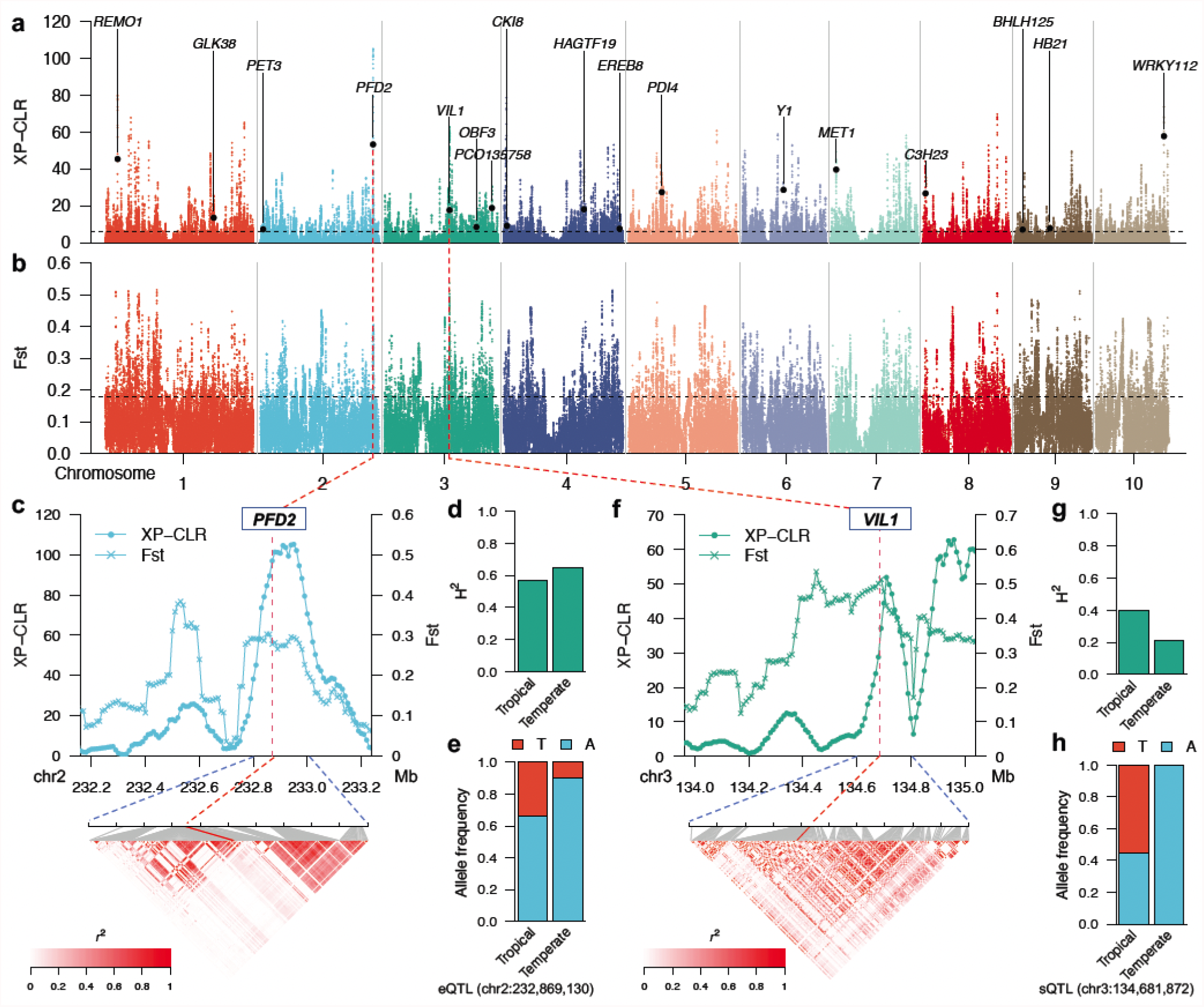
Genome-wide analysis of selective sweeps for temperate adaptation. (**a-b**) Genome-wide XP-CLR scanning (**a**) and fixation index (Fst) (**b**) in 100kb sliding windows with a 10kb step size. The dashed horizontal line indicates the top 10% threshold. Characterized genes with significant selective sweep signals that also showed reduction of expression heritability in temperate maize (except for *PFD2*) are indicated. (**c**) The maize homolog of Arabidopsis *PREFOLDIN-2* (*PFD2*) located near the highest XP-CLR peak shows a slight increase of expression heritability and selection of the A allele of its *cis*-eQTL in temperate maize. (**d**) *VIN3-LIKE PROTEIN-1* (*VIL1*) located near the highest Fst peak shows a reduction of expression heritability and strong selection of the A allele of its *cis*-sQTL in temperate maize.

### Involvement of transcriptional regulatory elements in phenotypic adaptation in temperate maize

We identified one or more significant trait-associated SNPs in a genome-wide analysis using data from 14 of 27 previously published organismal-level phenotypes related to flowering, development, disease susceptibility, and yield (Supplemental document 6), including the previously validated *Y1* for endosperm color (Figure 7a, Figure S6). After consolidating trait-associated SNPs into distinct peaks, we determined that candidate genes associated with organismal-level phenotypes and *cis*-regulatory elements with selection features for temperate adaptation such as reduced expression heritability in temperate maize or selective sweep signatures included candidate genes are potentially associated with endosperm color (Figure 7a & b), flowering time (Figure 7b & Figure S6), and kernel development (Figure 7c & Figure S6). *Y1* was previously reported to control maize endosperm color; we also identified *Y1* in this study as exhibiting both a selective sweep signature (Figure 7e; *first panel*) and reduced expression heritability (Figure 7e; *second panel*). Y1 expression was significantly higher (Figure 7e; *third panel*) in temperate maize and temperate maize inbreds predominantly carried the *G* allele at the *cis*-eQTL peak (Figure 7e; *fourth panel*). *Sugars will eventually be exported 2* (*SWEET2*, Zm00001d009365) was identified as a candidate gene associated with flowering time in this study (see Supplemental document 7) and exhibited selective sweep signatures (Figure 7 f; *first panel*) with a slightly increased expression heritability in temperate maize (Figure 7f; *second panel*). The *cis*-eQTLs and *cis*-sQTLs for *SWEET2* corresponded to two distinct XP-CLR peaks (Figure 7f; *first panel*). *SWEET2* expression was significantly higher in temperate maize relative to tropical maize (Figure 7f; *third panel*) while the *T* allele was less common in the temperate maize lines included in this study (Figure 7f; *fourth panel*). We also identified the DNA replication licensing factor gene *MCM4* (Zm00001d009374) as a candidate gene associated with flowering time whose gene region and eQTL were nearby XP-CLR peaks (Figure 7g; *first panel*). The expression heritability of *MCM4* was reduced by almost 50% in temperate maize compared to tropical maize (Figure 7g; *second panel*), while its median expression level decreased by 25% in temperate maize (Figure 7g; *third panel*). The *G* allele at the peak SNP for the associated *cis*-eQTL was much less common among temperate maize lines (Figure 7g; *fourth panel*). A gene encoding an aldolase superfamily protein (Zm00001d049559) was associated with upper kernel shape, with a *cis*-eQTL associated with its expression located within peaks for both XP-CLR and Fst (Figure 7h; *first panel*). The heritability of expression for this *Aldolase* was much lower in temperate maize than in tropical maize (Figure 7h; *second panel*) and exhibited a modest but significantly higher expression in temperate maize (Figure 7h; *third panel*). In agreement with the selective sweep signature of the *cis*-eQTL, the allele frequency for the peak SNP of this *cis*-eQTL differs substantially between tropical and temperate lines (Figure 7h; *fourth panel*). In addition, one of the SNPs linked to this gene was previously identified as a *cis*-eQTL at *Discolored 1* (*DSC1*, Zm00001d049872), which is associated with kernel development^63^. Both gene body and this eQTL mapped either within or nearby local Fst and XP-CLR peaks (Figure S8). *DSC1* exhibited significantly lower expression heritability in temperate maize compared to tropical maize, suggesting strong selection signals acting on its expression (Figure S8).

**Figure 7.**
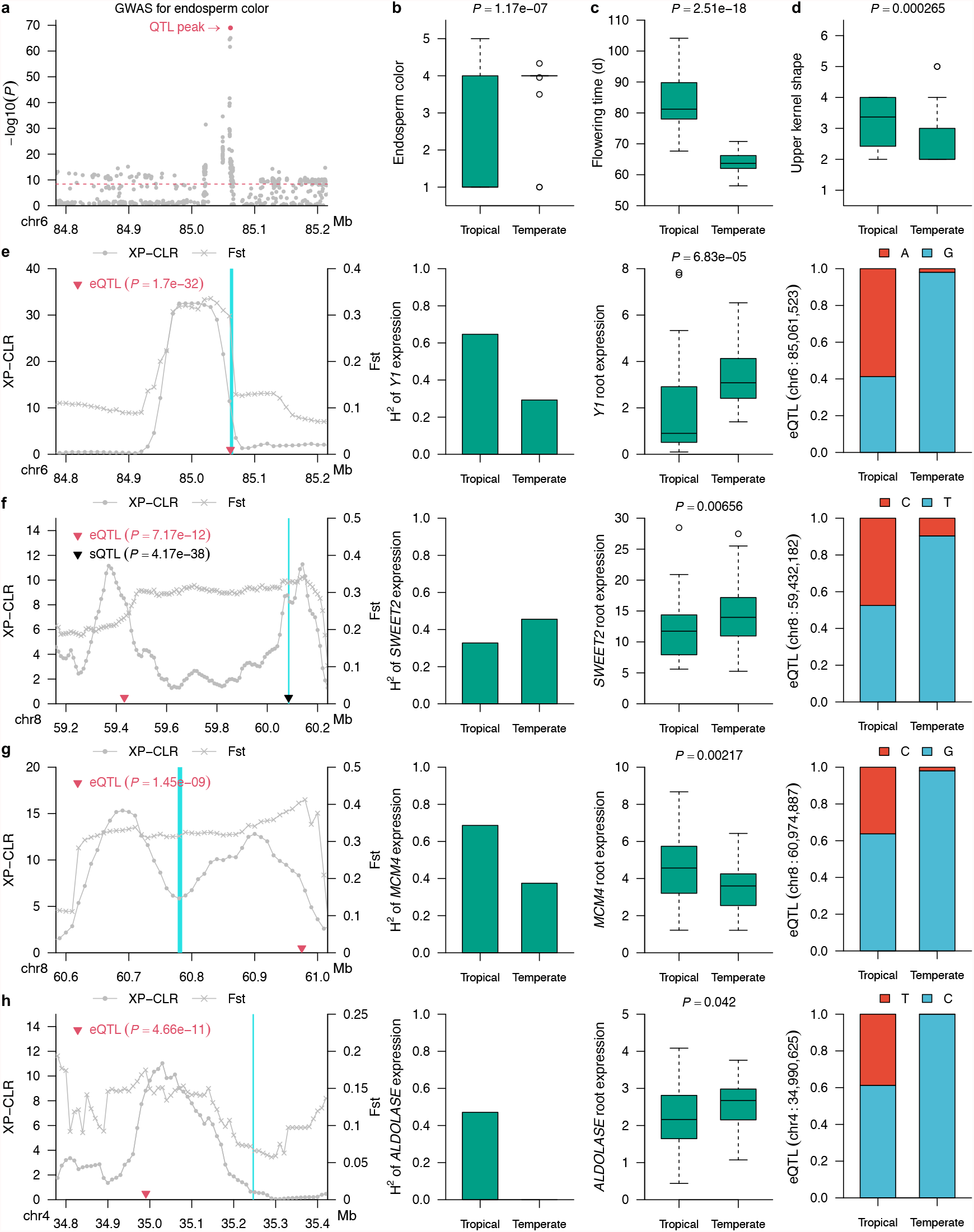
Candidate genes with differential selective sweeps for phenotypic adaptation in temperate maize. (**a**) Manhattan plot of GWAS for endosperm color in the population used in this study. The dashed horizontal line indicates the threshold, and the peak SNP is located in the *Y1* region. (**b**) Endosperm color in temperate maize compared to that in tropical maize. Trait is quantified using the code 1,white; 2,pale; 3,pale yellow; 4,yellow; 5, orange. (**c**) Flowering time (growing degree index from planting to 50% of anthesis in degrees Farenheit) is under selection in temperate maize compared to tropical maize. (**d**) Upper kernel shape is under selection in temperate maize compared to tropical maize. The trait is quantified using the code 1,shunken; 2,indented; 3,level; 4,rounded; 5,pointed; 6,strongly pointed. (**e-h**) From left to right, selective sweep signals (XP-CLR and Fst), expression broad sense heritability (H_2_), expression level (FPKM) in roots and *cis*-eQTL lead SNP allele frequency comparison between tropical and temperate maize for *Y1* (**e**), *SWEET2* (**f**), *MCM4* (**g**), and *ALDOLASE* (**h**).

## Discussion

The regulation of transcription and mRNA abundance is a critical intermediate step mediating how the genotype determines the phenotype and phenotypic plasticity in response to environmental variation. However, molecular phenotypes such as the abundance of individual transcripts are subject to differing degrees of genetic and non-genetic control, as are whole organism phenotypes. Here, we generated partially replicated data from a panel of maize lines originating from different parts of the globe. We estimated the extent of variation in transcript abundance explained by genetic factors with single-gene resolution by including biological replication of genetically identical individuals. In addition, by employing random fragmentation of cDNA molecules rather than targeted 3’ end sequencing of mRNAs, we quantified the fraction of variation in mRNA splicing that is genetically controlled.

We identified 1,272 genes with significantly lower broad sense heritability for their expression in temperate maize compared to tropical lines, indicating a transition to either become housekeeping genes or to be more environmentally responsive. The functions enriched in these genes included cell cycle regulation, DNA methylation and demethylation and mRNA binding activities, suggesting that the genetic regulation of gene expression and cell propagation are likely power houses driving adaptation to temperate environments from tropical maizes inbreds to temperate inbreds (Figure 1). In addition, the profiling of broad sense heritability by sequencing biological replicates allowed an estimation of all genetic factors (additive and non-additive) that explained variation in expression, in contrast to narrow sense heritability, which only measures additive genetic factors^64–67^. The inclusion of non-additive genetic factors is not trivial because non-additive factors such as dominance and epistasis explain a significant fraction of gene expression variation in both humans and plants^67,68^.

The eQTLs and sQTLs identified in this study did not capture all expression heritability due to the limited size of the population, which limited the statistical power to detect low-effect *trans*-eQTLs (Figure 2, S3). We conducted independent component analysis (ICA)^48^ and determined that the components are heritable with broad sense heritability ranging from zero to one, suggesting that these heritable components might represent certain co-expression modules that together exhibit detectable heritability when background noise from other heritable modules is removed. In humans, ICA has detected broad impact eQTLs when confounding factors within the expression matrix reduces the power of eQTL detection^69^. GWASs between the SNPs and these components also identified genomic regions harboring genes with global regulatory functions. For example, the peak on chromosome 4 at 79 Mb with enriched e-traits in the co-expression gene module was associated with independent component 32 (IC32), and the significant SNPs in this peak were strongly linked to Zm00001d050293, encoding trehalose-6-phosphate synthase 9 (ZmTRPS9). One of the mysteries surrounding trehalose metabolism is the high degree of redundancy between TRPSs and trehalose-6 phosphate phosphatases (TPPs or TRPPs). Although TRRS and TRPP homologs are expressed differently in response to the same stress in plants, in maize, the trehalose metabolic pathway is uniformly repressed while trehalose-6 phosphate levels are elevated, linking plant hormone signaling and trehalose metabolism^55^. In addition to the involvement of TRPS9 in the *tb1* mutant^55^, TRP9 was also reported to be involved in disease responses in tomato (*Solanum lycopersicum*) and southern rust^56,57^. TRPS catalyzes the formation of T6P; IC32 was associated with TRPS9 and was enriched for phytohormone signaling and plant pathogen interactions, providing another piece of evidence that trehalose metabolism is involved in plant hormone sensing and disease response. IC96 was associated with a large cluster of over 700 genes with a significant enrichment in phytohormone signaling, plant-pathogen interactions, and root hair development. IC96 was associated with a peak on chromosome 1 at 281 Mb corresponding to Zm00001d034004, encoding an Acyl-CoA N-acyltransferase with RING/FYVE/PHD-type zinc finger protein previously reported to be associated with plant height in response to drought stress in maize^58^. IC149 was associated with a SNP on chromosome 3 at 42.3 Mb which was linked to Zm00001d040415, encoding MIZU-KUSSEI 1 (MIZ1) that involves in lateral root development by maintaining auxin levels^52–54^. This finding is in agreement with cellulose and xylan biosynthesis reactions being enriched in the plant reactome for IC149 (Figure 4b). Due to the connection between a number of peaks and cluster functions identified by GWAS between SNPs and ICA components are reasonably supported by either literature or predictions. We concluded that the unknown or novel functions or connections we identified are due to a lack of current knowledge of the function of the candidate *trans*-acting regulators.

Rare alleles, mapping to *cis*-regulatory genic regions are usually deleterious, cause extremes in gene expression levels and are associated with lower seed fitness in maize^2^. We showed here that rare alleles in regions flanking *trans*-eQTLs are also significantly associated with extremes of gene expression (Figure 2e & f), suggesting that purging the rare alleles around *trans*-eQTLs might also be beneficial for fitness improvement in maize. In humans, spliceosome assembly co-opts RNA polymerase II for mRNA biosynthesis. Abolished spliceosome recruitment due to the artificial removal of introns leads to unprocessed RNA molecules that remain associated with RNA polymerase II, which eventually pauses on the nascent but unprocessed transcript^70^. The set of 666 genes with highly linked (co-localized) *cis*-eQTLs and sQTLs identified in this study may in fact support the possibility that a similar mechanism is at play in maize to regulate gene expression by eQTLs and sQTLs simultaneously (Figure 3). Furthermore, we also noticed that 33 of the 78 annotated genes were transcription factors and a positive correlation between expression levels and degree of splicing was observed in a significant portion of genes with intron retention variations would suggest a potential collaborative mode of action between transcription and splicing to achieve rapid response to environmental signals.

Maize domestication has reshaped the transcriptome; the genes with differential expression patterns were shown to be involved in biotic stress responses compared to teosinte (*Zea Mays ssp. Parvigliumis*)^71^. A significant portion of the genes previously identified to be targets for domestication and evolution by population genetics also exhibited altered expression patterns^71^. Temperate adaptation is a major part of maize domestication and largely contributed to the current global distribution of this crop. Liu et al^72^ attempted to dissect the genetic architecture of temperate adaptation of maize at both genomic and transcriptomic level, a set of 2700 differentially expressed genes involved in stress adaptation between temperate and tropical-subtropical maize lines^72^. Temperate regions constantly impose drought stress onto maize during the growing seasons^33^. Wang et al^73^ showed that a 366 bp insertion in the promoter of *ZmVPP1*, encoding a vacuolar-type H^+^ pyrophosphatase, controls the drought inducible expression of *ZmVPP1* to confer drought tolerance in maize^73^. In this study, we systematically investigated the involvement of expression regulatory elements in phenotypes associated with temperate adaptation, using a combination of genome wide eQTL and sQTL mapping, genome wide selective sweep detection among temperate and tropical maize subgroups included in the RNAseq population, and GWASs with phenotypes important for adaptation to temperate environments.

A set of 2,503 genes mapped within regions enriched for selective sweep signals, several of which are promising candidates for the phenotypic adaptation of temperate maize. For example, the maize homolog of Arabidopsis *VIL1* was previously shown to be associated with flowering time and showed the highest XP-CLR value of all tested genes^61,74^. Flowering time has long been considered one of the most important traits for temperate adaptation^32^. Further GWASs for 27 phenotypes, including endosperm color, flowering time, and kernel development, identified several associated candidate genes with *cis*-regulatory elements in selection signals (Figure 7). For all genes, either their *cis*-eQTLs, their *cis*-sQTLs, or both were within or close to the regions exhibiting strong selective sweep signals. For example, *Y1* encodes a phytoene synthetase involved in carotenoid biosynthesis^60^(Figure 7a & e). This gene is associated with endosperm color via GWASs (this study) and transcriptome wide association studies (TWAS)^75^ and this study (Figure 7a). Temperate maize was specifically selected for the *y1* allele, resulting in the greater accumulation of carotenoids in the endosperm compared to that of tropical maize^76^ (Figure 7b). We showed here that *ZmSWEET2* is associated with flowering time, and the upstream eQTL is nearby a local XP-CLR peak, in addition to one sQTL in the gene body near another selective sweep signal peak (Figure 7f). Its paralog ZmSWEET4c, was shown to be involved in seed filling, while ZmSWEET13s was also shown to be associated with flowering time^77^. The results in this study suggest *ZmSWEET2* as a promising candidate gene for flowering time regulation. We identified *MCM4* (*Minichromosome maintenance protein 4*) (Zm00001d009374) as another candidate gene for flowering time (Figure 7g). *MCM4* is a DNA helicase that forms a ring-shaped MCM complex along with other MCM subunits to activate DNA replication origin sites, followed by unwinding the DNA helix and formation of the DNA-replication fork^78–80^. After DNA replication is initiated, the MCM complex is released and prevented from reloading onto the nascent DNA^81,82^. The MCM complex is enriched in flowering bud tissue in Arabidopsis^80^. Based on the qTeller transcriptional database^83^, *ZnMCM4* is also stably expressed in maize flowering tissues (Figure S7). *BICELLULAR POLLEN1* (*BICE1*) in which mutations lead to defective gametogenesis in Arabidopsis, was shown to play a role in modulating DNA replication by interacting with MCM4^84^. Given that cell cycle is coupled with cell fate specification, these observations together strongly suggest roles for DNA replication in regulating flowering time. Zm00001d049559 encodes a transaldolase that we identified here as a candidate gene associated with kernel development with strong selection signals for expression and eQTLs (Figure 7h). This gene is an ortholog of Arabidopsis Clc-HYPERSENSITIVE MUTANT2 (GSM2)-like^85^. Arabidopsis GSM2 localizes to cotyledon chloroplast and contributes to scavenging reactive oxygen species in response to glucose during cotyledon development^85^. Transaldolases act one step upstream of transketolase to catalyze the oxidation of glucose-6-phosphate into ribulose-5-phosphate, which is a critical process in the oxidative pentose phosphate pathway (PPP). Defective PPP in chloroplasts is associated with lower oil and starch contents in the embryo^86^. Together, these data provide insights into one potential mode of action for *cis*-regulatory elements involved in maize temperate adaptation.

In conclusion, this study advances our understanding of how variation in gene expression may have supported the adaptation of maize to temperate environments. Our deployment of the ICA method in this study offered promising fundamental knowledge to explore potential latent *trans*-regulatory modules in the maize genome to capture missing heritability. Even though the genome assembly and annotation used in this study (the B73 genome version 4) represent the gold standard of a maize genome, structural variation such as presence/absence variation (PAV), copy number variation (CNV) and insertion/deletion (Indels) can affect the number of reads mapped to each genomic region. Future work should be therefore focus on pan-genome eQTL mapping to capture more gene expression heritability in maize.

## Materials and Methods

### Plant growth conditions

Plant tissue for this study was collected from maize plants grown between May 2017 and October 2019 in overlapping batches of 16 genotypes. Kernels were surface sterilized with chlorine gas in a desiccator. Surface sterilized kernels were hydrated in aerated 1 mM CaCl_2_ solution overnight before transfer to petri dishes containing paper towels soaked with 1 mM CaCl_2_. Petri dishes were sealed with micropore tape (3 M) and wrapped in black cloth before being placed in an incubator at 28-30 °C for 4-5 days to allow kernels to germinate.

Two of the successfully germinated kernels per genotype with radicle roots and coleoptiles of similar length were transferred to a semi-hydroponic growth system for vegetative development, consisting of a glass tube filled with 3 mm diameter glass beads and encased in PVC pipes to maintain dark conditions for the seedling root system^87^. All of the 16 PVCs pipes representing the 16 different maize lines were embedded in a rack platform connected to an intermittent watering system containing essential nutrients to support seedling growth (see Supplemental Table 3 for the recipe of the nutrient solution). The hydroponic growth system was placed in a growth chamber with 60% relative humidity, a 16-h-light/8-h-dark cycle, and 26°:8°day and night target temperatures.

Root tissue was harvested from the seedlings at 14 days of growth in the hydroponic system, between zeitgeber time 5 (ZT5) and ZT8 (with ZT0 being subjective dawn), and flash frozen in liquid nitrogen before storage at −80 °C until RNA extraction. Root tissue collection was conducted in a dark room solely illuminated by a bulb covered by a green filter (Cinegel #4490, Grand Stage Company, Chicago, IL).

### RNA extraction and sequencing

Frozen root samples were homogenized by grinding to a fine powder in liquid nitrogen. Total RNA was extracted with Trizol reagent from approximately 50 mg as per the manufacturer’s instructions. Total RNA was precipitated by centrifugation at 12,000 g for 15 min at 4°*C*. The resulting pellet was washed three times with 75% (v/v) ethanol before being resuspended in 40 µL of DEPC-treated water heated to 65°*C*.

RNA samples with RNA integrity number (RIN) <5 (Agilent 2100 Bioanalyzer) were discarded and new extractions conducted. RNA-seq libraries were constructed using the Illumina TruSeq v2 kit following the manufacturer’s published protocol^88^, pooled, and sequenced on an Illumina Nextseq 500 instrument with a target read length of 2×75 nucleotides and a target sequencing depth of 20 M paired-end reads per sample.

### Quantification of gene expression

The overall quality of RNAseq reads was assessed by FastQC^89^. Reads assigned to each RNA-seq library were filtered and quality trimmed using Trimmomatic (v 0.33) with parameter settings “-phred33 LEADING:3 TRAILING:3 slidingwindow:4:15 MINLEN:36 ILLUMINACLIP:TruSeq3-PE.fa:2:30:10”^90^. Trimmed reads were mapped to the B73_RefGen_v4 maize reference genome^91,92^ using STAR (v2.7) in two rounds^93^. In the first round, novel splice sites were identified from all libraries by mapping the reads to the reference genome with parameter settings “–alignIntronMin 20 –alignIntronMax 20000 –outSAMtype None –outSJfilterReads Unique –outSJfilterCountUniqueMin 10 3 3 3 –outSJfilterCountTotalMin 10 3 3 3”. In the second round, mapping of all libraries was conducted using an updated genome index including both annotated and newly identified splice sites from the first round of mapping with parameters “–alignIntronMin 20 –alignIntronMax 20000 –limitBAMsortRAM 5000000000 –outSAMstrandField intronMotif –alignSJoverhangMin 20 –outSAMtype BAM SortedByCoordinate”.

A mixture of in-house python scripts (deposited in the github repository associated with this article and the ‘prepDE.py’ python script provided by the StringTie (v2.1)^45^ package were employed to generate read counts. Estimated Fragments Per Kilobase of transcript per Million mapped reads (FPKM) values for all libraries were exported by Ballgown (v2.20.0)^94^.

### Genotype dataset preparation

Unimputed genotype calls from the set of 1,210 maize inbreds included in the maize hapmap3^40^ were filtered in a two step process. For the first step, line level filtering removed 588 lines where the missing data rate was ≥0.6 or the inbreeding coefficient was≤ 0.9. Of those lines that met the criteria for removal, 110 lines were retained, as they were included in the population employed in this study. In the second step, genetic markers with an observed minor allele frequency <0.01 among the remaining lines or a missing data rate >0.6 were removed. Missing data for markers that passed filtering in individuals were imputed using Beagle/5.1 with parameter settings “window=1 overlap=0.1 ne=1200”^95^. After imputation, sites where more than 20% of individuals were heterozygous were removed, resulting in a provisional marker set of 23,672,341 SNPs.

Individual RNA-seq samples were scored for each marker from the provisional marker set using GATK4 (v4.1) in GVCF mode^96^ using the binary sequence alignment format (BAM) files generated by STAR as described before. Individual libraries where >20% of genotyped markers were heterozygous were considered to represent contamination at either a genetic (pollen), sample (RNA isolation), or barcoding (library construction) level and were discarded from downstream analyses.

For samples representing genotypes not present in HapMap3 or samples where RNA-seq based SNP calling identified a set of markers discordant with published SNP data for the same line (<80% agreement at markers with minor allele frequencies >20%), downstream analyses employed SNP calls for the HapMap3 marker set generated using merged BAM files of all RNA-seq libraries for a given line and were imputed as described above. In all other cases, SNP data from whole genome resequencing in HapMap3 were employed. The final SNP genotype dataset employed for downstream analyses resulted from filtering the provisional marker set to remove markers with >2% heterozygous SNP calls or a minor allele frequency <0.05 among the set of genotypes employed in this study. These filtering criteria resulted in a set of 12,191,984 markers employed for all downstream analyses.

A kinship matrix was calculated using 244,683 independent SNP markers (LD R^2^ ≤0.2) using PLINK (v1.9)^97^. Principal components of genetic variation were calculated using a randomly selected set of 1,000,000 markers from the total set of markers using tassel/5.2. LD heatmaps were generated using the R package Gaston^98^.

### Gene expression heritability

Broad sense heritability for the expression of individual genes was estimated as the total variation explained by genotype (sigmal_G) as a proportion of total variation (sigma_P) 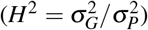. sigma_G and sigma_P were estimated using the lme4 R package and data for 219 replicated genotypes, or separately for a set of 22 out of 40 (22 with replicates) genotypes of high confidence tropical origin and 39 out of 52 (39 with replicates) genotypes of high confidence temperate origin^38^, fitting genotype as a random effect^99^. All other unexplained variations were considered as random errors.

To test for significant shifts in the heritability of genes in the same functional groups (same GO terms) in the whole RNA-seq population (340 genotypes), a two-sample Kolmogorov-Smirnov test was performed using the heritability of genes assigned the same GO annotations published as part of the Maize-GAMER dataset^100^ and the heritability of other genes assigned to the other GO terms. For testing purposes, the population set was defined as 19,565 genes with FPKM ≥ 1 in more than 80% of all RNAseq samples included in this study. The resulting *p* values for GO terms were corrected for FDR <0.05 (Benjamini Hochberg method of multiple testing correction), and the median heritability of the genes assigned to the test GO terms with a ≥20% difference compared to the background population was considered significant. For the purpose of visualization, the redundancy of significant GO terms identified by this method was reduced by REVIGO with the default method (SimRel) and mall similarity (0.5)^101^. The original list of GO terms is provided in Supplemental Table S2

For genes with expression heritabilities reduced over 80% in temperate maize compared to those in tropical maize, GO term enrichment among these genes was performed by goatools with the background set defined as the 19,565 genes described before^102^. *p* values for each GO terms were corrected for a FDR <0.05 (Benjamini Hochberg method of multiple testing correction), and GO term redundancy was reduced in REVIGO with the same settings as above^101^ with further manual optimization for visualization. The original list of GO terms is provided in Supplemental Table S4.

### eQTL mapping unique peak calling

eQTL mapping was conducted using MatrixEQTL(v2.3)^39^ with gene expression values (FPKM) first normalized via the Box-Cox transformation^103^ and with five principle components of population structure included as covariates. For each gene, eQTLs were classified as *cis* if they were located within 1 Mb upstream or downstream of the gene’s annotated transcription start site or transcription stop site. *p*-values for each gene-SNP pair were adjusted using Bonferroni correction (*p* = 4.1e^−9^)^104^. Peak calling was conducted by merging trait associated SNPs located within 1 Mb of each other. Peaks with at least 3 significantly SNPs were retained for downstream analyses.

### Quantification of RNA splicing and splicing-QTL mapping

Introns used in this analysis were assembled by StringTie (v2.1)^45^ based on the reads mapped to the B73_RefGen_V4 reference genome using STAR^93^ (as described above). PSI (Percent Spliced-in) for each intron was used to quantify RNA splicing events. The formula used to calculate PSI was ucount/(depth/size), where ucount is the number of uniquely mapped junction reads for an intron using the iexpr function in Ballgown (v2.20.0)^94^, depth is the sum of uniquely mapped read depths per base for the intron calculated using samtools bedcov function (-Q 255)^105^, and size is the length of the intron in base pairs. Only splicing events (introns) with five or more unique junction reads in at least 5% of all RNAseq samples were kept for PSI calculation. PSI was set to missing if depth was zero. Introns with a standard deviation >0.1 and a missing data rate <5% of PSI value in the population were used for splicing QTL mapping using the same methods described above for eQTL mapping. Following the methods for splicing QTL mapping proposed by Chen et al^20^, only significant hits with allele affects (beta) higher than 0.05 were kept for peak consolidation using the method described above.

### Analysis of selective sweep genes under regional domestication

The fixation index (Fst) values were calculated from the SNP dataset for the 40 maize genotypes with high-confident tropical origins and 52 maize genotypes with high-confidence temperate origins using VCFtools (v0.1.16) with a window size of 100 kb and a step size of 10 kb^106^. Using the same set of SNP, whole-genome cross-population composite likelihood ratio (XP-CLR) scores were calculated using xpclr (v1.0)^107^ to compare tropical and temperate populations using parameters “-w1 0.0005 100 100 1 -p1 0.95” for each chromosome following previous studies with modifications^108–110^. The genetic position of each SNP was inferred from a published genetic map constructed for the nested association mapping (NAM) population of US inbred lines assuming uniform rate of recombination between mapped markers^111^. To enable comparisons between Fst and XP-CLR scores, XP-CLR scores were averaged within each of the 100 kb Fst window^109^. For both Fst and XP-CLR, the top 10% of 100 kb windows were considered highly differentiated. Following a previously proposed QC protocol, windows with ratios of nucleotide diversity (*π*) (tropical/temperate) lower than the genome wide average were removed from the sets of both Fst and XP-CLR highly differentiated regions^109,110^. A gene was considered a candidate target of selection for temperate adaptation if it was located entirely within windows identified as highly differentiated by both Fst and XP-CLR analyses. The R package Gaston was used to visualize LD (linkage disequilibrium) heatmaps in the candidate gene regions^98^.

### Organismal phenotype GWAS

A set of 27 trait datasets with data for *geq* 250 maize genotypes for which RNA-seq data was generated as part of this study were obtained from USDA GRIN (https://www.ars-grin.gov/) or Peiffer et al.^112^. Raw trait values were Log transformed prior to analyses. GWAS was conducted using the algorithm GEMMA (v0.98.3) with a kinship matrix and the first five principal components of population structure included as covariates^113^. Genes considered potentially causal were: 1) those where either a trait associated SNP or a SNP in >0.8 LD with a trait-associated SNP was located within 50 kb of the annotated start and stop position of the gene; or 2) those where an eQTL identified for expression level of the gene was also identified as a trait associated SNP and in >0.8 LD with a trait associated SNP.

### Gene models identified by independent component analysis

Independent component analysis was conducted with the FastICA algorithm as implemented in the FastICA package (version 1.2-2)^114,115^. Based on singular value decomposition (SVD), 164 components explained 80% of the total variance in the expression matrix of 19,565 gene models used for eQTL mapping with broad sense heritabilties ≥ 0.05. ICA was conducted using parameter settings “nbComp” = 164, “maximum iteration” = 500 and the default function “logcosh”. Only the 43 components with kurtosis values >6 and at least 10 genes associated with the module at an FDR threshold of 0.001 were kept for further analysis.

The coefficients of these 43 independent components assigned to each individual were used to calculate the Best Linear Unbiased Predictions (BLUPs) using the lme4 package^99^. BLUPs for each principal component were Log transformed before being subjected to GWAS analysis using the same method as the organismal phenotype GWAS described above. Peaks were consolidated using the same method as described above. Peaks supported by fewer than three significant SNPs were excluded from downstream analysis. Candidate causal genes for each peak of interest were identified based on their relative location and LD (R^2^ >0.8) with the SNPs within the peak regions. Plant reactome enrichment was performed using ShinyGO (v0.66)^116^ with the set of 19,565 genes used for eQTL mapping defined as the background gene set.

## Data availability statement

RNAseq data for root tissues of 572 maize genotypes are available at NCBI under the BioProject: PRJNA793045. All of the in-house code for analysis are accessible at github repository : https://github.com/gsun2unl/eGWAS

## Author’s contribution

J. C. S., G. S., and H. H., conceived, designed and directed the study. P. W., M. L. G., developed the plant growing workflow and collected root samples. R. V. M., collected W., and L. B., profiled primary metabolites. C. C., L. S., Y. Y., C. D., K. B., and R. O directed and performed paspalum genome sequencing. J. S., C.P., and J. J., directed and performed paspalum genome assembly. K. D., P. Q., and T. G., constructed genetic maps used for genome assembly. S. S., performed the annotation of the genome assembly. B. Y., and B. Z., designed and conducted the quantification of ATG8 and ATG8-PE. C. Z., A.L. and H. Y assisted with the RNAseq analysis. B. S., designed and conducted flow cytometry experiments. J. C. S., and G. S., drafted the manuscript. The final version of the manuscript was generated with input and contributions from N. W., S. S., J. J., B. Z., P. Q., H. Y., C. Z., K. D., B. S., B. Y., T. O., J. S., All authors approved the final version of the manuscript.

## Acknowledgements

This work was supported by a National Science Foundation Award (OIA-1557417) to JCS, CZ, KVD, & DPS. This project was completed utilizing the Holland Computing Center of the University of Nebraska, which receives support from the Nebraska Research Initiative. We thank Bert Devilbiss for his technical assistance and organization and Allyn Pella for organizing the seed increases.

---

**Table S1.** Chi-square test for significance of association between cis-elements and selection of expressed genes in maize

|  | With cis-elements | Without cis-elements | Sum |
| --- | --- | --- | --- |
| Selected | 634 | 424 | 1058 |
| Not selected | 11908 | 6599 | 18607 |
| Sum | 12542 | 7023 | 19565 |
| p value =0.003961452236186191 |  |  |  |

**Table S2.** Peaks enriched for etrait in the corresponding co-expression modules

| leadSNP | eTraitNumber | ModuleSize | EnrichmentFactor | chi2_Pval | IC |
| --- | --- | --- | --- | --- | --- |
| 1:111188891(IC46) | 10 | 144 | 4.104775092312857 | 4.5345650349964485e-06 | IC46 |
| 3:42334086(IC149) | 26 | 207 | 2.3835495766504073 | 4.895733220520719e-06 | IC149 |
| 4:79115907(IC32) | 58 | 283 | 1.6714414280390681 | 3.0395172767548555e-05 | IC32 |
| 1:280872052(IC96) | 45 | 743 | 1.8779074851812907 | 1.3287397712783146e-05 | IC96 |
| 3:163218947(IC96) | 23 | 743 | 3.3834973721211754 | 6.645470949158957e-10 | IC96 |
| 4:242527467(IC96) | 30 | 743 | 2.2128097537803817 | 8.146261296758717e-06 | IC96 |
| 10:34688181(IC96) | 145 | 743 | 2.666343037812883 | 2.1797782090934917e-38 | IC96 |

**Figure S1.**
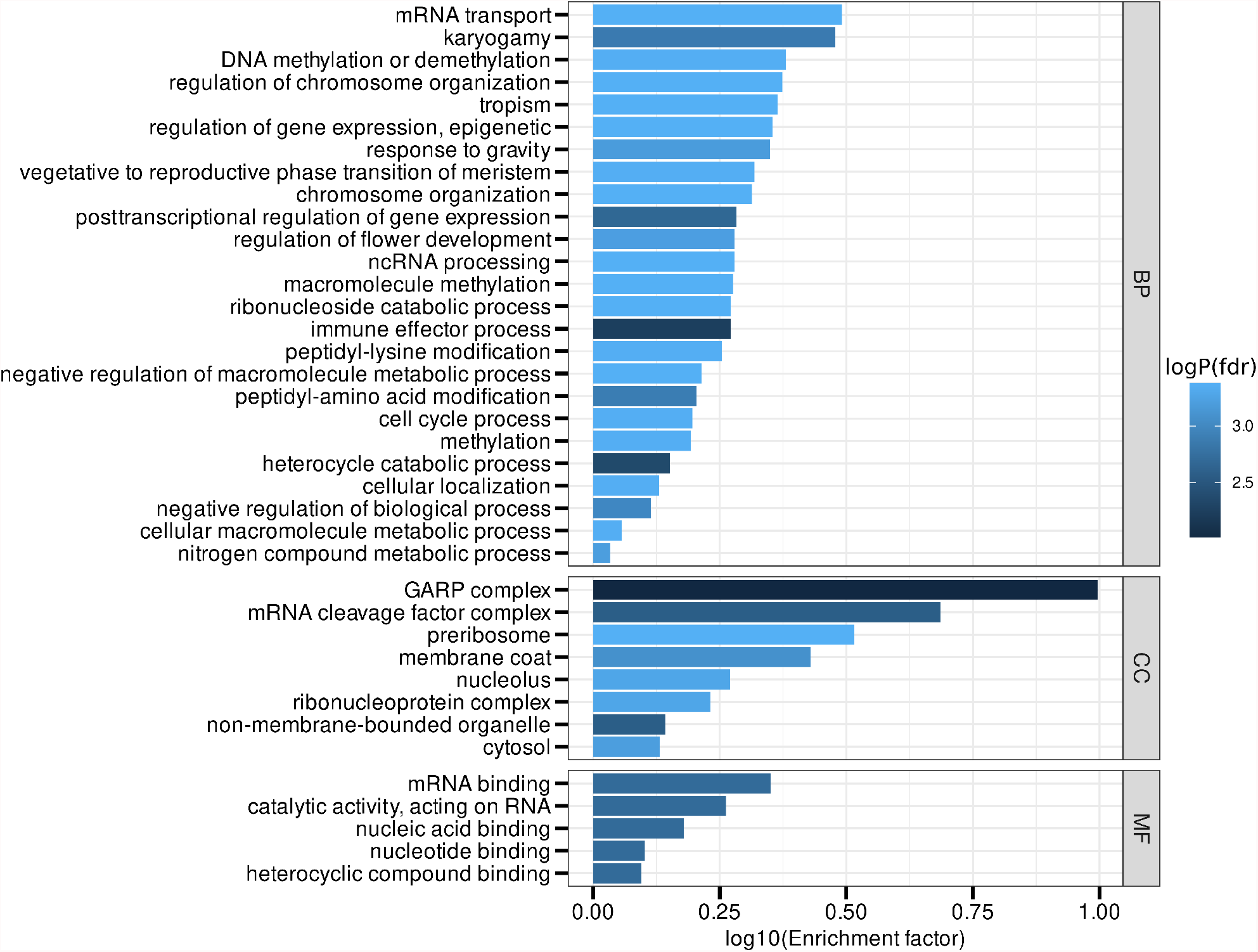
Gene Ontology terms enriched in the genes exhibiting reduced expression heritability in temperate maize. Significantly enriched GO terms (FDR ≤0.05) for 1,272 genes exhibiting at least an 80% reduction of expression heritability in the temperate maize population relative to the tropical maize population. Bars indicate the Log_10_-transformed enrichment factor (number of genes associated with the overrepresented GO term in the study gene set over the number of genes associated with the GO term in the background gene set). The background gene set is the 19,565 genes used for the eGWAS. Negative Log_10_-transformed multi-test corrected *p*-values are color-coded.

**Figure S2.**
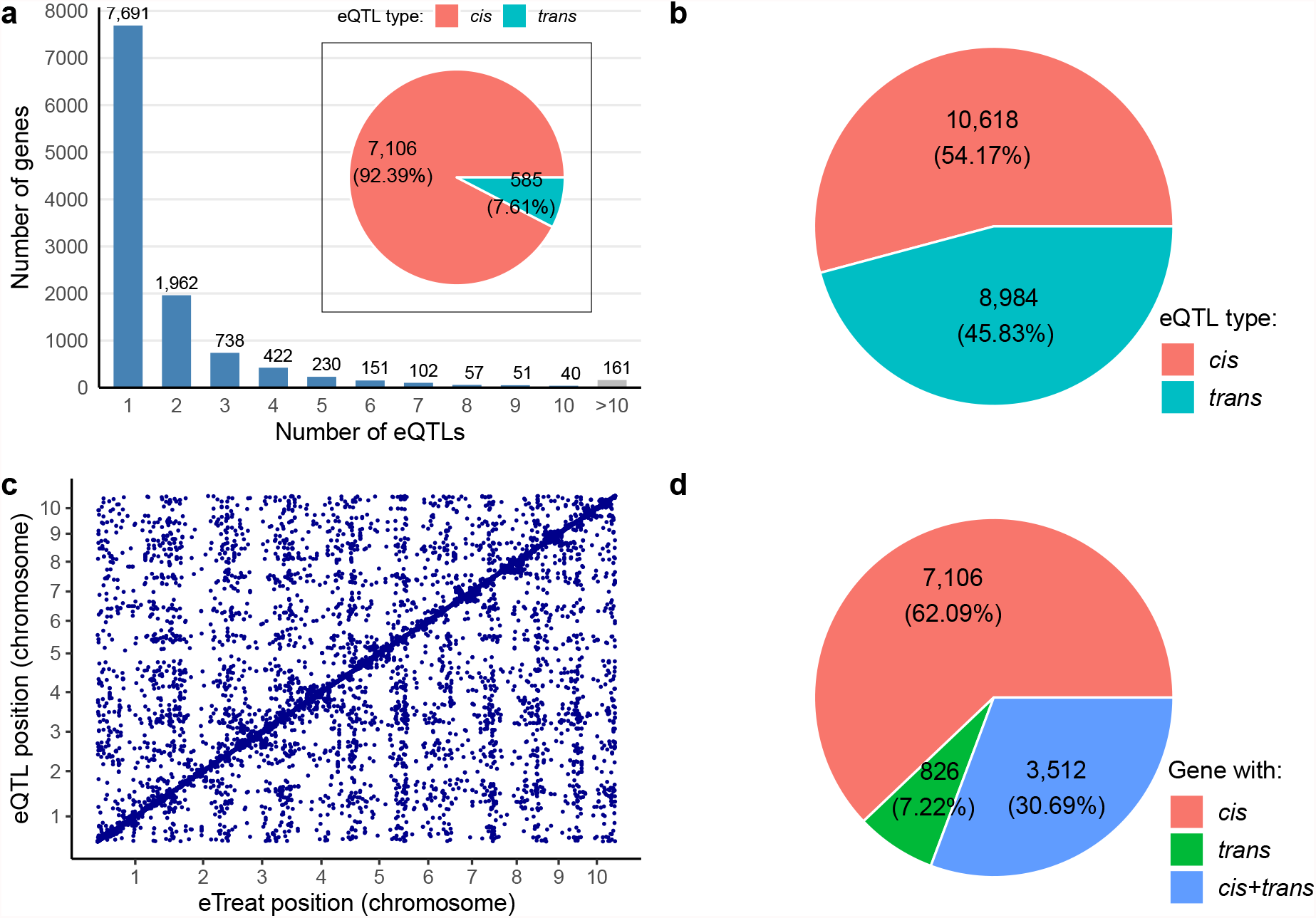
eQTL composition and distribution. (**a**) Number of genes with one or more significant peaks and the proportion of *cis*-and *trans*-eQTLs in genes with only one peak. (**b**) The proportion of *cis*-and *trans*-eQTLs identified by eQTL mapping in this study. (**c**) Positional correlation of e-traits and eQTLs; the dots along the diagonal indicate *cis*-eQTLs, while other dots indicate *trans*-eQTLs. (**d**) Proportions among the 11,444 e-traits with *cis*-or *trans*-or both eQTLs.

**Figure S3.**
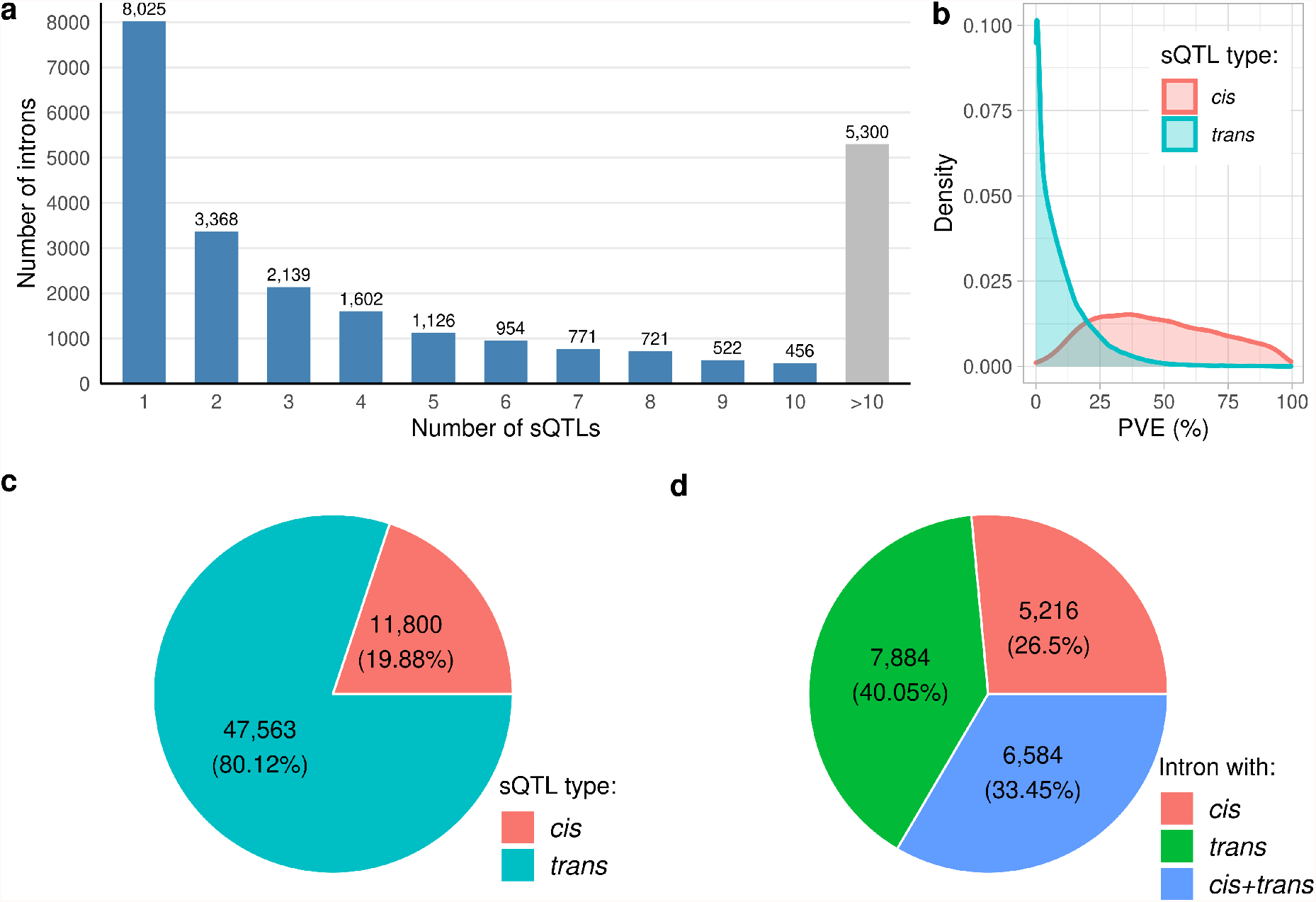
sQTL composition and distribution. (**a**) Number of introns with one or more significant sQTL peak. (**b**) Expression variation explained by their associated *cis*-and *trans*-eQTLs. (**c**) Proportion of *cis*-and *trans*-sQTLs identified by sQTL mapping in this study. (**d**) Proportions among the 19,684 splicing events associated with just *cis*-, just *trans*-, or both *cis*- and *trans*-sQTLs.

**Figure S4.**
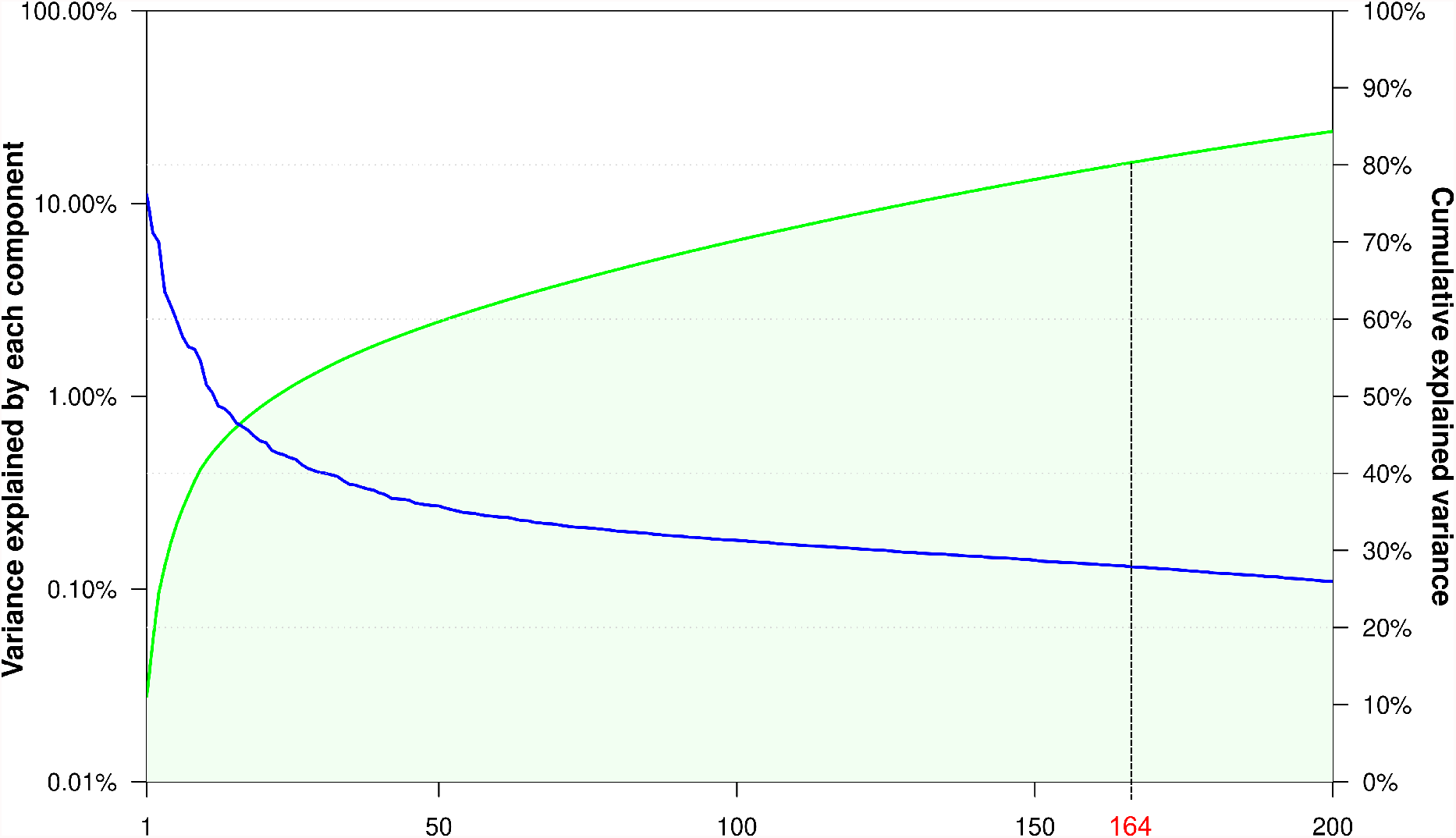
Cumulative percentage of variance explained by independent components.

**Figure S5.**
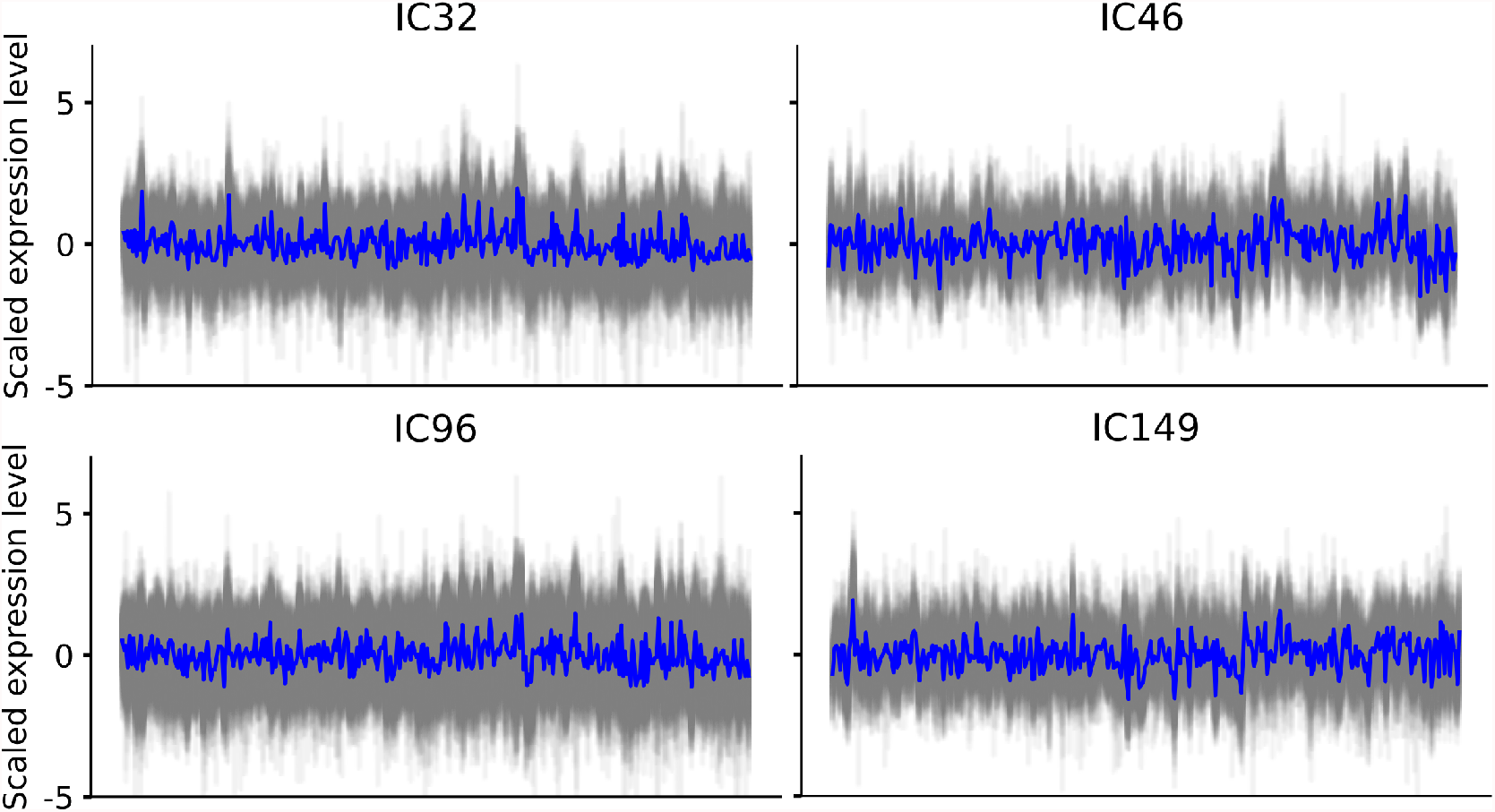
Expression patterns of the genes within in each of the co-expression modules associated with IC32, IC46, IC96 and IC146.

**Figure S6.**
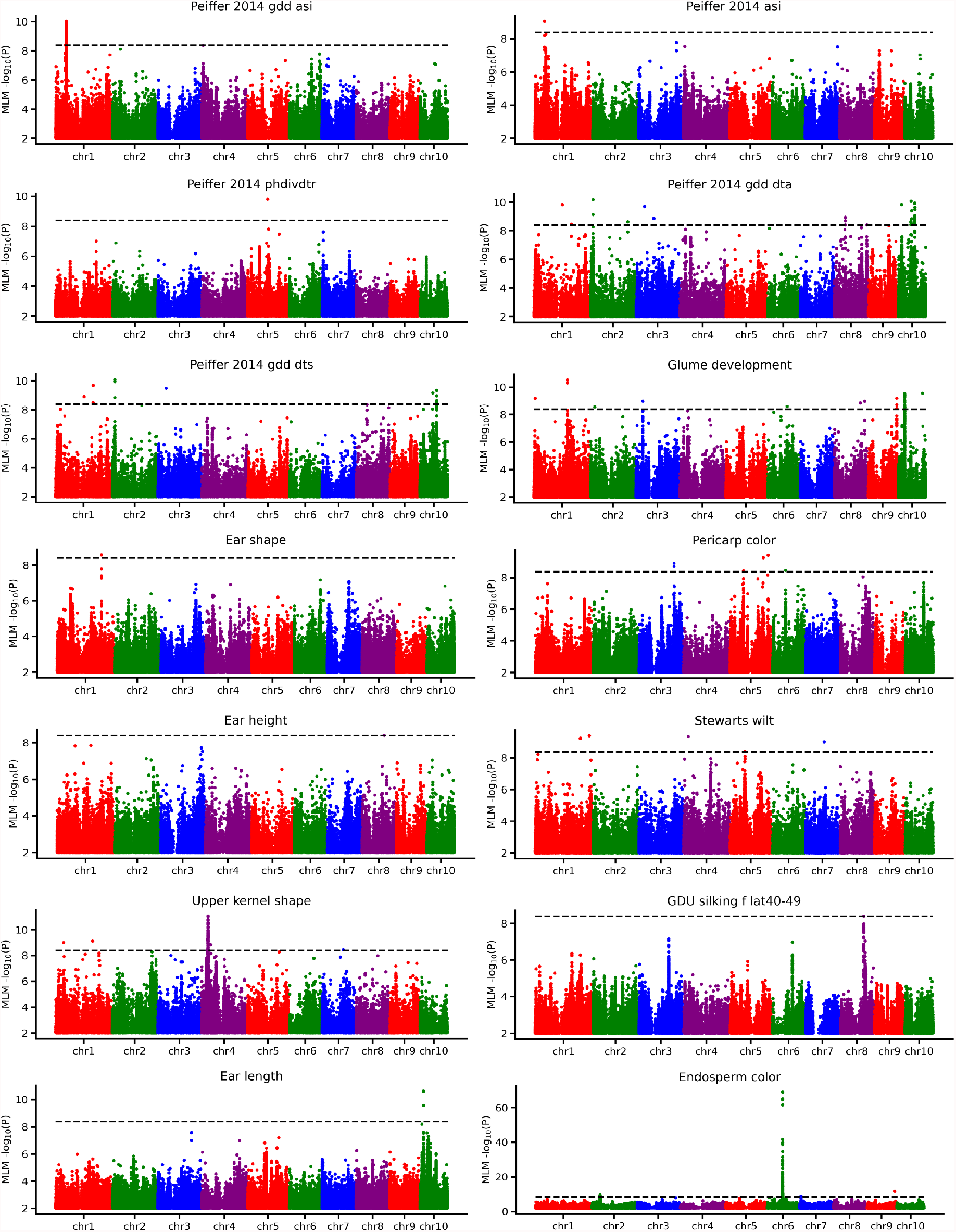
GWAS for 14 phenotypes with significant signals. *p*-values of the association between each of the 12,191,984 SNPs across the genome and the phenotypic variation for the 14 phenotypes across the 340 genotypes are shown.

**Figure S7.**
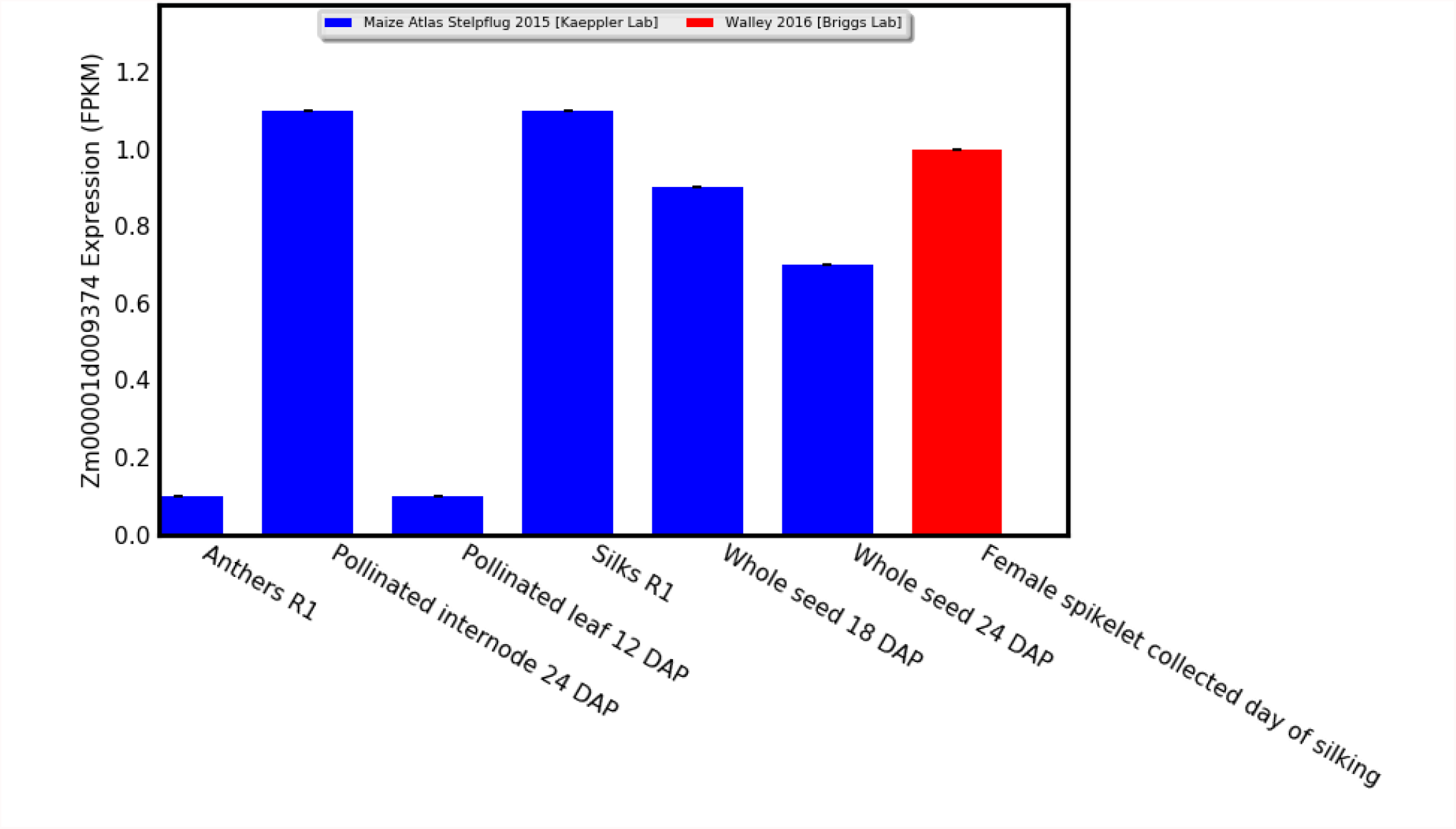
Expression patterns (FPKM) of *MCM4* across different flowering tissues in maize reference inbred B73.

**Figure S8.**
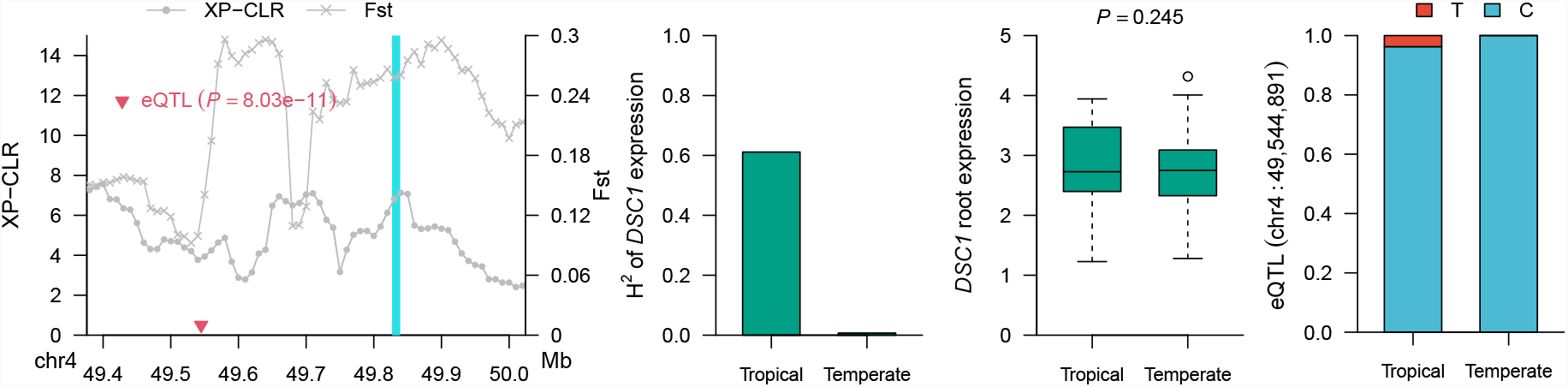
Selection features of *DSC1* associated with kernel development. From left to right, selective sweep signals (XP-CLR and Fst), expression broad sense heritability (H^2^), expression level (FPKM) in roots, and *cis*-eQTL lead SNP allele frequency comparison between tropical and temperate maize

## References

1. Unterseer, S. et al. A comprehensive study of the genomic differentiation between temperate dent and flint maize. Genome biology 17, 1–14 (2016).

2. Kremling, K. A. et al. Dysregulation of expression correlates with rare-allele burden and fitness loss in maize. Nature 555, 520–523 (2018).

3. Rodgers-Melnick, E., Vera, D. L., Bass, H. W. & Buckler, E. S. Open chromatin reveals the functional maize genome. Proc. Natl. Acad. Sci. 113, E3177–E3184 (2016).

4. Brem, R. B., Yvert, G., Clinton, R. & Kruglyak, L. Genetic dissection of transcriptional regulation in budding yeast. Science 296, 752–755 (2002).

5. DeCook, R., Lall, S., Nettleton, D. & Howell, S. H. Genetic regulation of gene expression during shoot development in arabidopsis. Genetics 172, 1155–1164 (2006).

6. Swanson-Wagner, R. A. et al. Paternal dominance of trans-eqtl influences gene expression patterns in maize hybrids. science 326, 1118–1120 (2009).

7. West, M. A. et al. Global eqtl mapping reveals the complex genetic architecture of transcript-level variation in arabidopsis. Genetics 175, 1441–1450 (2007).

8. Li, L. et al. Mendelian and non-mendelian regulation of gene expression in maize. PLoS genetics 9, e1003202 (2013).

9. Liu, H. et al. Distant eqtls and non-coding sequences play critical roles in regulating gene expression and quantitative trait variation in maize. Mol. Plant 10, 414–426 (2017).

10. Christie, N. et al. Systems genetics reveals a transcriptional network associated with susceptibility in the maize–grey leaf spot pathosystem. The Plant J. 89, 746–763 (2017).

11. Wang, X. et al. Genome-wide analysis of transcriptional variability in a large maize-teosinte population. Mol. Plant 11, 443–459 (2018).

12. Tu, X. et al. Reconstructing the maize leaf regulatory network using chip-seq data of 104 transcription factors. Nat. communications 11, 1–13 (2020).

13. Gore, M. A. et al. A first-generation haplotype map of maize. Science 326, 1115–1117 (2009).

14. Tibbs Cortes, L., Zhang, Z. & Yu, J. Status and prospects of genome-wide association studies in plants. The Plant Genome 14, e20077 (2021).

15. Liu, S. et al. Mapping regulatory variants controlling gene expression in drought response and tolerance in maize. Genome biology 21, 1–22 (2020).

16. Fu, J. et al. Rna sequencing reveals the complex regulatory network in the maize kernel. Nat. communications 4, 1–12 (2013).

17. Pang, J. et al. Kernel size-related genes revealed by an integrated eqtl analysis during early maize kernel development. The Plant J. 98, 19–32 (2019).

18. Lohman, B. K., Weber, J. N. & Bolnick, D. I. Evaluation of tagseq, a reliable low-cost alternative for rna seq. Mol. ecology resources 16, 1315–1321 (2016).

19. Thatcher, S. R. et al. Genome-wide analysis of alternative splicing in zea mays: landscape and genetic regulation. The Plant Cell 26, 3472–3487 (2014).

20. Chen, Q. et al. Genome-wide association analyses reveal the importance of alternative splicing in diversifying gene function and regulating phenotypic variation in maize. The Plant Cell 30, 1404–1423 (2018).

21. Yu, J. et al. Alternative splicing of os lg 3b controls grain length and yield in japonica rice. Plant Biotechnol. J. 16, 1667–1678 (2018).

22. Chen, M. et al. Fine mapping of a major qtl for flag leaf width in rice, qflw4, which might be caused by alternative splicing of nal1. Plant cell reports 31, 863–872 (2012).

23. Zhang, Z. & Xiao, B. Comparative alternative splicing analysis of two contrasting rice cultivars under drought stress and association of differential splicing genes with drought response qtls. Euphytica 214, 1–16 (2018).

24. Yu, H. et al. Genome-wide discovery of natural variation in pre-mrna splicing and prioritizing causal alternative splicing to salt stress response in rice. New Phytol. (2021).

25. Kesari, R. et al. Intron-mediated alternative splicing of arabidopsis p5cs1 and its association with natural variation in proline and climate adaptation. Proc. Natl. Acad. Sci. 109, 9197–9202 (2012).

26. Cubillos, F. A., Coustham, V. & Loudet, O. Lessons from eqtl mapping studies: non-coding regions and their role behind natural phenotypic variation in plants. Curr. opinion plant biology 15, 192–198 (2012).

27. Arnaud, N., Lawrenson, T., Østergaard, L. & Sablowski, R. The same regulatory point mutation changed seed-dispersal structures in evolution and domestication. Curr. Biol. 21, 1215–1219 (2011).

28. Konishi, S. et al. An snp caused loss of seed shattering during rice domestication. Science 312, 1392–1396 (2006).

29. Studer, A., Zhao, Q., Ross-Ibarra, J. & Doebley, J. Identification of a functional transposon insertion in the maize domestication gene tb1. Nat. genetics 43, 1160–1163 (2011).

30. Matsuoka, Y. et al. A single domestication for maize shown by multilocus microsatellite genotyping. Proc. Natl. Acad. Sci. 99, 6080–6084 (2002).

31. Swarts, K. et al. Genomic estimation of complex traits reveals ancient maize adaptation to temperate north america. Science 357, 512–515 (2017).

32. Buckler, E. S. et al. The genetic architecture of maize flowering time. Science 325, 714–718 (2009).

33. Camus-Kulandaivelu, L. et al. Maize adaptation to temperate climate: relationship between population structure and polymorphism in the dwarf8 gene. Genetics 172, 2449–2463 (2006).

34. Tollenaar, M. & Wu, J. Yield improvement in temperate maize is attributable to greater stress tolerance. Crop. science 39, 1597–1604 (1999).

35. Revilla, P. et al. Association mapping for cold tolerance in two large maize inbred panels. BMC plant biology 16, 1–10 (2016).

36. Salvi, S. et al. Conserved noncoding genomic sequences associated with a flowering-time quantitative trait locus in maize. Proc. Natl. Acad. Sci. 104, 11376–11381 (2007).

37. Flint-Garcia, S. A. et al. Maize association population: a high-resolution platform for quantitative trait locus dissection. The plant journal 44, 1054–1064 (2005).

38. Romay, M. C. et al. Comprehensive genotyping of the usa national maize inbred seed bank. Genome biology 14, R55 (2013).

39. Shabalin, A. A. Matrix eqtl: ultra fast eqtl analysis via large matrix operations. Bioinformatics 28, 1353–1358 (2012).

40. Bukowski, R. et al. Construction of the third-generation zea mays haplotype map. Gigascience 7, gix134 (2018).

41. Liu, B., Gloudemans, M. J., Rao, A. S., Ingelsson, E. & Montgomery, S. B. Abundant associations with gene expression complicate gwas follow-up. Nat. genetics 51, 768–769 (2019).

42. Wilkinson, M. E., Charenton, C. & Nagai, K. Rna splicing by the spliceosome. Annu. review biochemistry 89 (2020).

43. Li, S. et al. Global co-transcriptional splicing in arabidopsis and the correlation with splicing regulation in mature rnas. Mol. plant 13, 266–277 (2020).

44. Jia, J. et al. Post-transcriptional splicing of nascent rna contributes to widespread intron retention in plants. Nat. Plants 6, 780–788 (2020).

45. Pertea, M. et al. Stringtie enables improved reconstruction of a transcriptome from rna-seq reads. Nat. biotechnology 33, 290–295 (2015).

46. Shaul, O. How introns enhance gene expression. The international journal biochemistry & cell biology 91, 145–155 (2017).

47. Bommert, P., Je, B. I., Goldshmidt, A. & Jackson, D. The maize gα gene compact plant2 functions in clavata signalling to control shoot meristem size. Nature 502, 555–558 (2013).

48. Rotival, M. et al. Integrating genome-wide genetic variations and monocyte expression data reveals trans-regulated gene modules in humans. PLoS genetics 7, e1002367 (2011).

49. Li, H. & Adali, T. A class of complex ica algorithms based on the kurtosis cost function. IEEE Transactions on Neural Networks 19, 408–420 (2008).

50. Naithani, S. et al. Plant reactome: a resource for plant pathways and comparative analysis. Nucleic acids research gkw932 (2016).

51. Naithani, S. et al. Plant reactome: a knowledgebase and resource for comparative pathway analysis. Nucleic acids research 48, D1093–D1103 (2020).

52. Moriwaki, T. et al. Hormonal regulation of lateral root development in arabidopsis modulated by miz1 and requirement of gnom activity for miz1 function. Plant Physiol. 157, 1209–1220 (2011).

53. Yamazaki, T. et al. Miz1, an essential protein for root hydrotropism, is associated with the cytoplasmic face of the endoplasmic reticulum membrane in arabidopsis root cells. FEBS letters 586, 398–402 (2012).

54. Kobayashi, A. et al. A gene essential for hydrotropism in roots. Proc. Natl. Acad. Sci. 104, 4724–4729 (2007).

55. Dong, Z. et al. The regulatory landscape of a core maize domestication module controlling bud dormancy and growth repression. Nat. communications 10, 1–15 (2019).

56. Zhang, H. et al. Virus-induced gene silencing-based functional analyses revealed the involvement of several putative trehalose-6-phosphate synthase/phosphatase genes in disease resistance against botrytis cinerea and pseudomonas syringae pv. tomato dc3000 in tomato. Front. plant science 7, 1176 (2016).

57. Sun, G., Mural, R. V., Turkus, J. D. & Schnable, J. C. Quantitative resistance loci to southern rust mapped in a temperate maize diversity panel. Phytopathology PHYTO–04 (2021).

58. Li, C. et al. Numerous genetic loci identified for drought tolerance in the maize nested association mapping populations. BMC genomics 17, 1–11 (2016).

59. Zhao, D.-Z., Wang, G.-F., Speal, B. & Ma, H. The excess microsporocytes1 gene encodes a putative leucine-rich repeat receptor protein kinase that controls somatic and reproductive cell fates in the arabidopsis anther. Genes & Dev. 16, 2021–2031 (2002).

60. Buckner, B., Kelson, T. L. & Robertson, D. S. Cloning of the y1 locus of maize, a gene involved in the biosynthesis of carotenoids. The Plant Cell 2, 867–876 (1990).

61. Sung, S., Schmitz, R. J. & Amasino, R. M. A phd finger protein involved in both the vernalization and photoperiod pathways in arabidopsis. Genes & development 20, 3244–3248 (2006).

62. Zhang, X. & Qi, Y. Genetic architecture affecting maize agronomic traits identified by variance heterogeneity association mapping. Genomics (2021).

63. Takacs, E. M., Suzuki, M. & Scanlon, M. J. Discolored1 (dsc1) is an adp-ribosylation factor-gtpase activating protein required to maintain differentiation of maize kernel structures. Front. plant science 3, 115 (2012).

64. Condon, A. & Richards, R. Broad sense heritability and genotype ×environment interaction for carbon isotope discrimination in field-grown wheat. Aust. J. Agric. Res. 43, 921–934 (1992).

65. Petretto, E. et al. Heritability and tissue specificity of expression quantitative trait loci. PLoS genetics 2, e172 (2006).

66. Albert, F. W., Bloom, J. S., Siegel, J., Day, L. & Kruglyak, L. Genetics of trans-regulatory variation in gene expression. Elife 7, e35471 (2018).

67. Li, Z., Gao, N., Martini, J. W. & Simianer, H. Integrating gene expression data into genomic prediction. Front. Genetics 10, 126 (2019).

68. Brown, A. A. et al. Genetic interactions affecting human gene expression identified by variance association mapping. Elife 3, e01381 (2014).

69. Ju, J. H., Shenoy, S. A., Crystal, R. G. & Mezey, J. G. An independent component analysis confounding factor correction framework for identifying broad impact expression quantitative trait loci. PLoS computational biology 13, e1005537 (2017).

70. Martins, S. B. et al. Spliceosome assembly is coupled to rna polymerase ii dynamics at the 3’ end of human genes. Nat. structural & molecular biology 18, 1115–1123 (2011).

71. Swanson-Wagner, R. et al. Reshaping of the maize transcriptome by domestication. Proc. Natl. Acad. Sci. 109, 11878–11883 (2012).

72. Liu, H. et al. Genomic, transcriptomic, and phenomic variation reveals the complex adaptation of modern maize breeding. Mol. plant 8, 871–884 (2015).

73. Wang, X. et al. Genetic variation in zmvpp1 contributes to drought tolerance in maize seedlings. Nat. genetics 48, 1233–1241 (2016).

74. Sung, S., Schmitz, R. J. & Amasino, R. The role of vin3-like genes in environmentally induced epigenetic regulation of flowering. Plant signaling & behavior 2, 127–128 (2007).

75. Li, D., Liu, Q. & Schnable, P. S. Twas results are complementary to and less affected by linkage disequilibrium than gwas. Plant Physiol. (2021).

76. Palaisa, K., Morgante, M., Tingey, S. & Rafalski, A. Long-range patterns of diversity and linkage disequilibrium surrounding the maize y1 gene are indicative of an asymmetric selective sweep. Proc. Natl. Acad. Sci. 101, 9885–9890 (2004).

77. Bezrutczyk, M. et al. Impaired phloem loading in zmsweet13a, b, c sucrose transporter triple knock-out mutants in zea mays. New Phytol. 218, 594–603 (2018).

78. Gambus, A. et al. Gins maintains association of cdc45 with mcm in replisome progression complexes at eukaryotic dna replication forks. Nat. cell biology 8, 358–366 (2006).

79. Liu, W., Pucci, B., Rossi, M., Pisani, F. M. & Ladenstein, R. Structural analysis of the sulfolobus solfataricus mcm protein n-terminal domain. Nucleic acids research 36, 3235–3243 (2008).

80. Shultz, R. W., Lee, T.-J., Allen, G. C., Thompson, W. F. & Hanley-Bowdoin, L. Dynamic localization of the dna replication proteins mcm5 and mcm7 in plants. Plant physiology 150, 658–669 (2009).

81. Namdar, M. & Kearsey, S. E. Analysis of mcm2–7 chromatin binding during anaphase and in the transition to quiescence in fission yeast. Exp. cell research 312, 3360–3369 (2006).

82. Kearsey, S. E. & Labib, K. Mcm proteins: evolution, properties, and role in dna replication. Biochimica et Biophys. Acta (BBA)-Gene Struct. Expr. 1398, 113–136 (1998).

83. Woodhouse, M. R. et al. qteller: A tool for comparative multi-genomic gene expression analysis. Bioinformatics (2021).

84. Long, Y.-P., Xie, D.-J., Zhao, Y.-Y., Shi, D.-Q. & Yang, W.-C. Bicellular pollen 1 is a modulator of dna replication and pollen development in arabidopsis. New Phytol. 222, 588–603 (2019).

85. Zheng, M., Zhu, C., Yang, T., Qian, J. & Hsu, Y.-F. Gsm2, a transaldolase, contributes to reactive oxygen species homeostasis in arabidopsis. Plant molecular biology 104, 39–53 (2020).

86. Spielbauer, G. et al. Chloroplast-localized 6-phosphogluconate dehydrogenase is critical for maize endosperm starch accumulation. J. experimental botany 64, 2231–2242 (2013).

87. Lopez-Guerrero, M. G. et al. A glass bead-semi hydroponic system for intact maize root exudate analysis and phenotyping. Plant Methods (in press).

88. Schroeder, A. et al. The rin: an rna integrity number for assigning integrity values to rna measurements. BMC molecular biology 7, 1–14 (2006).

89. Andrews, S. et al. Fastqc: a quality control tool for high throughput sequence data (2010).

90. Bolger, A. M., Lohse, M. & Usadel, B. Trimmomatic: a flexible trimmer for illumina sequence data. Bioinformatics 30, 2114–2120 (2014).

91. Schnable, P. S. et al. The b73 maize genome: complexity, diversity, and dynamics. science 326, 1112–1115 (2009).

92. Jiao, Y. et al. Improved maize reference genome with single-molecule technologies. Nature 546, 524–527 (2017).

93. Dobin, A. et al. Star: ultrafast universal rna-seq aligner. Bioinformatics 29, 15–21 (2013).

94. Frazee, A. C. et al. Ballgown bridges the gap between transcriptome assembly and expression analysis. Nat. biotechnology 33, 243–246 (2015).

95. Browning, B. L., Zhou, Y. & Browning, S. R. A one-penny imputed genome from next-generation reference panels. The Am. J. Hum. Genet. 103, 338–348 (2018).

96. Poplin, R. et al. Scaling accurate genetic variant discovery to tens of thousands of samples. BioRxiv 201178 (2017).

97. Chang, C. C. et al. Second-generation plink: rising to the challenge of larger and richer datasets. Gigascience 4, s13742–015 (2015).

98. Hervé Perdry, D. B., Claire Dandine-Roulland & Kettner, L. gaston: Genetic Data Handling (QC, GRM, LD, P.A. & Linear Mixed Models. R package version 1.5.7. Université Paris-Saclay (2020).

99. Bates, D., Sarkar, D., Bates, M. D. & Matrix, L. The lme4 package. R package version 2, 74 (2007).

100. Wimalanathan, K., Friedberg, I., Andorf, C. M. & Lawrence-Dill, C. J. Maize go annotation—methods, evaluation, and review (maize-gamer). Plant Direct 2, e00052 (2018).

101. Supek, F., Bošnjak, M., škunca, N. & šmuc, T. Revigo summarizes and visualizes long lists of gene ontology terms. PloS one 6, e21800 (2011).

102. Klopfenstein, D. et al. Goatools: A python library for gene ontology analyses. Sci. reports 8, 1–17 (2018).

103. Osborne, J. Improving your data transformations: Applying the box-cox transformation. Pract. Assessment, Res. Eval. 15, 12 (2010).

104. Vialatte, F.-B. & Cichocki, A. Split-test bonferroni correction for qeeg statistical maps. Biol. Cybern. 98, 295–303 (2008).

105. Li, H. et al. The sequence alignment/map format and samtools. Bioinformatics 25, 2078–2079 (2009).

106. Danecek, P. et al. The variant call format and vcftools. Bioinformatics 27, 2156–2158 (2011).

107. Chen, H., Patterson, N. & Reich, D. Population differentiation as a test for selective sweeps. Genome research 20, 393–402 (2010).

108. Hufford, M. B. et al. Comparative population genomics of maize domestication and improvement. Nat. genetics 44, 808–811 (2012).

109. Su, T. et al. A genomic variation map provides insights into the genetic basis of spring chinese cabbage (brassica rapa ssp. pekinensis) selection. Mol. plant 11, 1360–1376 (2018).

110. Wang, B. et al. Genome-wide selection and genetic improvement during modern maize breeding. Nat. genetics 52, 565–571 (2020).

111. Li, C. et al. Construction of high-quality recombination maps with low-coverage genomic sequencing for joint linkage analysis in maize. BMC biology 13, 1–12 (2015).

112. Peiffer, J. A. et al. The genetic architecture of maize height. Genetics 196, 1337–1356 (2014).

113. Zhou, X. & Stephens, M. Genome-wide efficient mixed-model analysis for association studies. Nat. genetics 44, 821 (2012).

114. Hyvarinen, A. Fast and robust fixed-point algorithms for independent component analysis. IEEE transactions on Neural Networks 10, 626–634 (1999).

115. Marchini, J., Heaton, C., Ripley, B. & Ripley, M. B. The fastica package (2007).

116. Ge, S. X., Jung, D. & Yao, R. Shinygo: a graphical gene-set enrichment tool for animals and plants. Bioinformatics 36, 2628–2629 (2020).

